# Divergent short-term plasticity creates parallel pathways for computation and behavior in an olfactory circuit

**DOI:** 10.1101/2024.04.29.591740

**Authors:** Hyong S. Kim, James M. Jeanne

## Abstract

To enable diverse sensory processing and behavior, central circuits use divergent connectivity to create parallel pathways. However, linking subcellular mechanisms to the circuit-level segregation of computation has been challenging. Here, we investigate the generation of parallel processing within a divergent network in the *Drosophila* olfactory system, where single projection neurons target multiple types of lateral horn neuron (LHN). One LHN type generates sustained responses and adapts divisively to encode temporal odor contrast. The other generates transient responses and adapts subtractively to encode a form of positive temporal prediction error. These coding differences originate from subcellular differences in short-term plasticity in projection neuron axons. Prediction error arises from strongly facilitating synapses, which depend on the presynaptic priming factor Unc13B. The temporal contrast code arises from mildly depressing synapses that engage additional gain control implemented by the Na^+^/K^+^ ATPase in the postsynaptic neuron. Each LHN type makes corresponding dynamic contributions to behavioral odor attraction. Subcellular synaptic specialization is a compact and efficient way to generate diverse parallel information streams.

## INTRODUCTION

Many stages of sensory processing in the brain are characterized by a limited number of neuron types that expand onto larger and more diverse downstream populations. This architecture allows for greater specialization later in sensory processing^1^. For instance, specialized transient and sustained representations emerge downstream of receptor neurons in both vision and olfaction^2-4^.

The elemental unit of expansion is divergence, where the axon of a single presynaptic neuron connects to multiple postsynaptic targets. Even within that one axon, specialization can already begin, when presynaptic short-term plasticity differs for different target neurons^5-11^. Although this subcellular specification of plasticity is widespread throughout the brain, we do not know how it implements different computation and behavior.

Addressing this issue involves three questions. First, how does sensory coding diversify downstream of a single neuron? Second, what are the cellular and subcellular molecular differences that produce this diversity? Finally, how do the different downstream neurons contribute to behavioral function?

To answer these questions, one would ideally identify individual neurons that form distinct synapse types onto different target neurons, directly compare their sensory coding properties *in vivo*, and genetically perturb them while measuring physiology and behavior. A useful experimental system for this purpose is the *Drosophila* lateral horn, a compact olfactory network with stereotyped synaptic connections^12-15^. Antennal lobe projection neurons (PNs) gather input from peripheral olfactory receptor neurons (ORNs), reliably encoding temporal odor patterns and sending their outputs to the lateral horn^16-18^. There, each PN axon diverges to form excitatory synaptic connections onto dozens of anatomically distinct types of lateral horn neuron (LHN)^3,12-14,19,20^, some of which instruct behavioral attraction or aversion^21,22^.

We aimed to understand how divergence from PNs to LHNs creates parallel channels for computation and behavior. We identify two LHN types that receive input from the same PNs yet differ in the transience of their responses to odor. One generates sustained responses, faithfully tracks odor dynamics, and adapts divisively to encode temporal odor contrast. The other generates transient responses, tracks only low frequency odor dynamics, and adapts subtractively to encode a form of positive temporal prediction error. We then trace the mechanisms that enable this divergent functionality, starting from target-neuron specific short-term plasticity in a single PN axon. Ultimately, this divergence contributes to separate control of sustained and transient behavioral responses to odor. Our results provide a mechanistic understanding of how an elemental motif of neuronal architecture – divergence – creates temporally diverse computation.

## RESULTS

In this study, we used sequences of odor pulses to drive time-locked packets of spikes in presynaptic PNs and monitored how different postsynaptic LHNs transform them and contribute to dynamic behavior (**Figure 1A**). The parallel computations performed by divergent circuits like this one may be most amenable to mechanistic study when activity is isolated to individual presynaptic neurons, so in our physiology experiments we focused on low concentration “private” odor stimuli that drive only one PN at a time.

**Figure 1:**
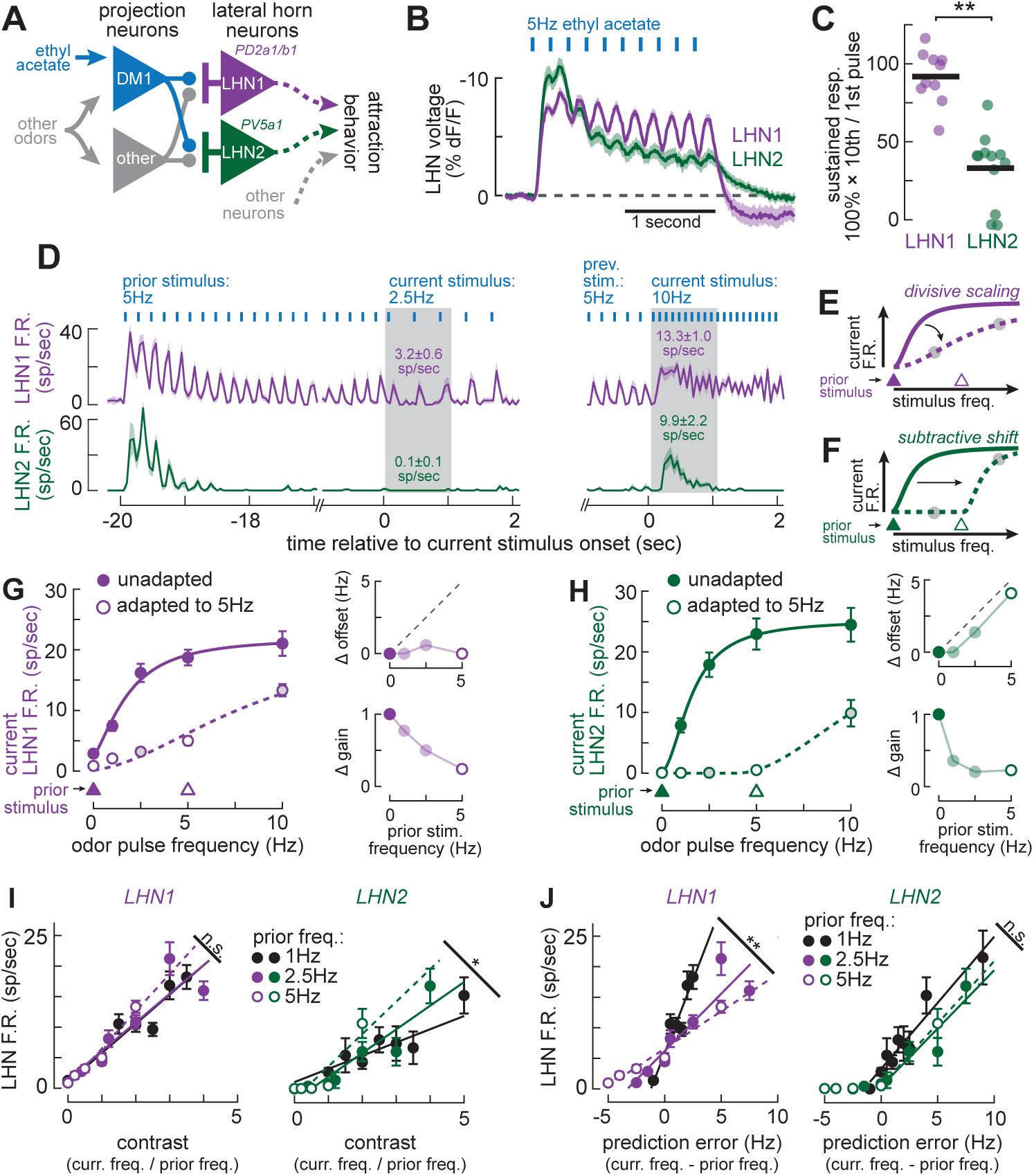
Distinct adaptation creates parallel representations of contrast and prediction error. **A)** Schematic of the microcircuit investigated in this study. Low concentration ethyl acetate activates DM1 without activating other PNs. LHN1 and LHN2 receive common inputs from DM1, as well as several other common PN types. Both LHN types contribute to attraction behavior, which also depends on other neuron types. **B)** Mean (± s.e.m.) voltage change ArcLight fluorescence) of LHN1 (n = 10) and LHN2 (n = 12) in response to identical stimuli. Voltage scale is inverted because fluorescence decreases with voltage. **C)** Comparison of sustained responsiveness (10th pulse amplitude as a percentage of 1st pulse amplitude) in LHN1 and LHN2. T test: ** p < 0.001. **D)** Mean (± s.e.m.) peristimulus time histograms (PSTHs) of spike rates in LHN1 (n = 10) and LHN2 (n = 7). The stimulus is ethyl acetate pulsed at 5Hz for 20 seconds prior to switching to either 2.5Hz (left) or 10Hz (right). Shaded areas denote 1-second analysis window for computing mean adapted spike rates (corresponding to shaded circles in panels G and H). **(E)** Schematic of adaptive divisive modulation. With a strong prior stimulus, the current firing rate curve scales along its input axis, reducing gain. Triangles at bottom denote strength of prior stimulus. **(F)** Schematic of adaptive subtractive modulation. With a strong prior stimulus, the current firing rate curve shifts along its input axis, generating an offset. Triangles at bottom denote strength of prior stimulus. **(G)** Left: mean (± s.e.m.) ethyl acetate responses for LHN1 (n = 9-16) for a range of stimulus frequencies without adaptation (solid circles) and adapted to 5Hz (open circles). Shaded circles correspond to shaded regions in (D). Curve was fit to unadapted (prior) spike rates, and then only offset and gain were adjusted to fit the adapted data (STAR Methods). Prior pulse frequencies are plotted in triangles at bottom. Right: offset and gain adjustments for the best fits to each adaptation frequency. **(H)** Same as (G) but for LHN2 (n = 6-16). **(I)** LHN1 (left) and LHN2 (right) spike rates as a function of temporal contrast (current pulse frequency divided by prior frequency). Best linear fits for each prior frequency have indistinguishable slopes for LHN1 but not LHN2, indicating that LHN1 encodes contrast better. * ANCOVA interaction, p = 0.011. **(J)** LHN1 (left) and LHN2 (right) spike rates as a function of temporal prediction error (prior pulse frequency subtracted from current frequency). Best linear fits for each prior frequency have indistinguishable slopes for LHN2 but not LHN1, indicating that LHN2 encodes positive prediction error (negative prediction errors do not yield positive spike rates). ** ANCOVA interaction, p = 0.005.

We targeted the DM1 PN because it has good genetic access^23^, it can be privately activated by ethyl acetate^24^, and it contributes to food attraction behavior^25,26^. We focused on two LHN types, called “PD2a1/b1” and “PV5a1” because DM1 makes strong anatomical connections onto both^12,14^ (**Figure S1A,B**) and both contribute to attraction behavior^21,27^. For simplicity, we refer to these LHN types as “LHN1” and “LHN2,” respectively.

In pilot experiments, we identified ethyl acetate delivery parameters that allowed varying levels of DM1 activity while remaining private (**Figure S1C-G**; **STAR Methods**). Because near-threshold odors can be difficult to control, we focused on the highest concentration of ethyl acetate that we found to be private and instead varied the odor pulse frequency (keeping pulse duration fixed). Higher frequencies delivered more odor in a fixed time window and led to increasing DM1 spike rates, without recruiting other glomeruli. Wind causes odor plumes to separate into discrete filaments, so the odor pulses likely resemble patterns that would occur naturally^28^.

### LHNs form sustained and transient representations

Voltage imaging from LHN1 and LHN2 revealed that they both entrain their voltages to 5Hz odor pulses, but they adapted differently (**Figure 1B**). LHN1 sustained large depolarizations over ten pulses, dropping only ∼10%. LHN2’s response was much more transient, dropping by ∼70% after ten pulses (**Figure 1C**). These different magnitudes of adaptation imply that each LHN type forms different temporal representations of the same dynamic odors.

To characterize these representations, we next made patch-clamp recordings from LHN1 and LHN2. We delivered 5Hz odor pulses for 20 seconds to drive responses into steady state and then either increased or decreased the pulse frequency (**Figure 1D**). LHN1 spike rates dropped over time, but still faithfully tracked each odor pulse in steady state and during increases or decreases in stimulation frequency. LHN2 spikes completely adapted away in steady state, and only responded transiently to increases in stimulation frequency. Notably, transient LHN2 activity did not require synaptic inhibition (**Figure S2A-C**). The distinction in transience did not require pulsed stimulation or private odor concentrations (**Figure S2D-H**). Distinct transience was also observed for a different odor, methyl acetate (**Figure S2I,J**), which privately activates the DM4 PN (and also targets both LHN types **Figure S1**). Thus, these LHNs generate distinct dynamics across a range of odor dynamics, concentrations, and identities.

### Distinct adaptation creates parallel representations of contrast and prediction error

Adaptive modulation of sensory tuning curves can be broadly classified as divisive (i.e. reducing gain; **Figure 1E**), subtractive (a rightward offset; **Figure 1F**), or a combination of both. To characterize these properties, we measured LHN responses to a range of odor pulse frequencies (0-10Hz), with or without prior stimulation (1-5Hz; we refer to this prior stimulation interchangeably as “adaptation”). Without prior stimulation, both LHN1 and LHN2 spike rates (averaged over the first 1 second of stimulation) followed similar saturating functions of pulse frequency, similar to other olfactory neurons^24^ (**Figure 1G,H**, **STAR Methods**). With prior stimulation, LHN1 reduced its gain, depending on prior stimulus frequency (**Figure 1G**). LHN2 both reduced its gain and generated an offset, but only the offset depended on the prior stimulus frequency (**Figure 1H**).

Divisive modulation (i.e., stretching the x-axis) enables spike rates to encode temporal contrast: the ratio of the current stimulus to the prior stimulus^29^. Accordingly, LHN1 spike rates corresponded better to contrast coding than did LHN2 (**Figure 1I**). Subtractive modulation (i.e., shifting the x-axis) enables only the difference between the current and prior stimulus to be encoded, a simple form of a positive prediction error (where the prediction is that prior stimuli will continue unchanged)^30^. Accordingly, LHN2 spike rates corresponded better to prediction error coding than LHN1 (**Figure 1J**).

In short, the two LHNs represent the same stimuli in different ways. LHN1 exhibits the hallmarks of contrast coding, with a sustained responses that faithfully track variable pulse frequencies. LHN2 exhibits the hallmarks of positive prediction error coding, with transient responses that only yield spikes for unexpectedly strong stimuli.

### LHN adaptation specializes upon pre-adapted PN input

As the sole source of odor-evoked excitation to LHN1 and LHN2 under our stimulus conditions, DM1 is an information processing bottleneck. Implementing common components of adaptation in DM1 would be a useful strategy to alleviate some of the burden on LHNs. Of course, any preprocessing must necessarily preserve all the information needed by both LHN types.

We thus measured the transience of DM1 spike rate responses to 5Hz odor pulses. Over several seconds, DM1 activity declined, indicating that some adaptation is already present at this stage of processing, as expected^31-33^ (**Figure 2A**). However, responses to each pulse declined by only ∼30% in DM1, compared to ∼75% in LHN1 and ∼100% in LHN2 (**Figure 2B**). Thus, DM1 provides LHNs with preprocessed inputs.

**Figure 2:**
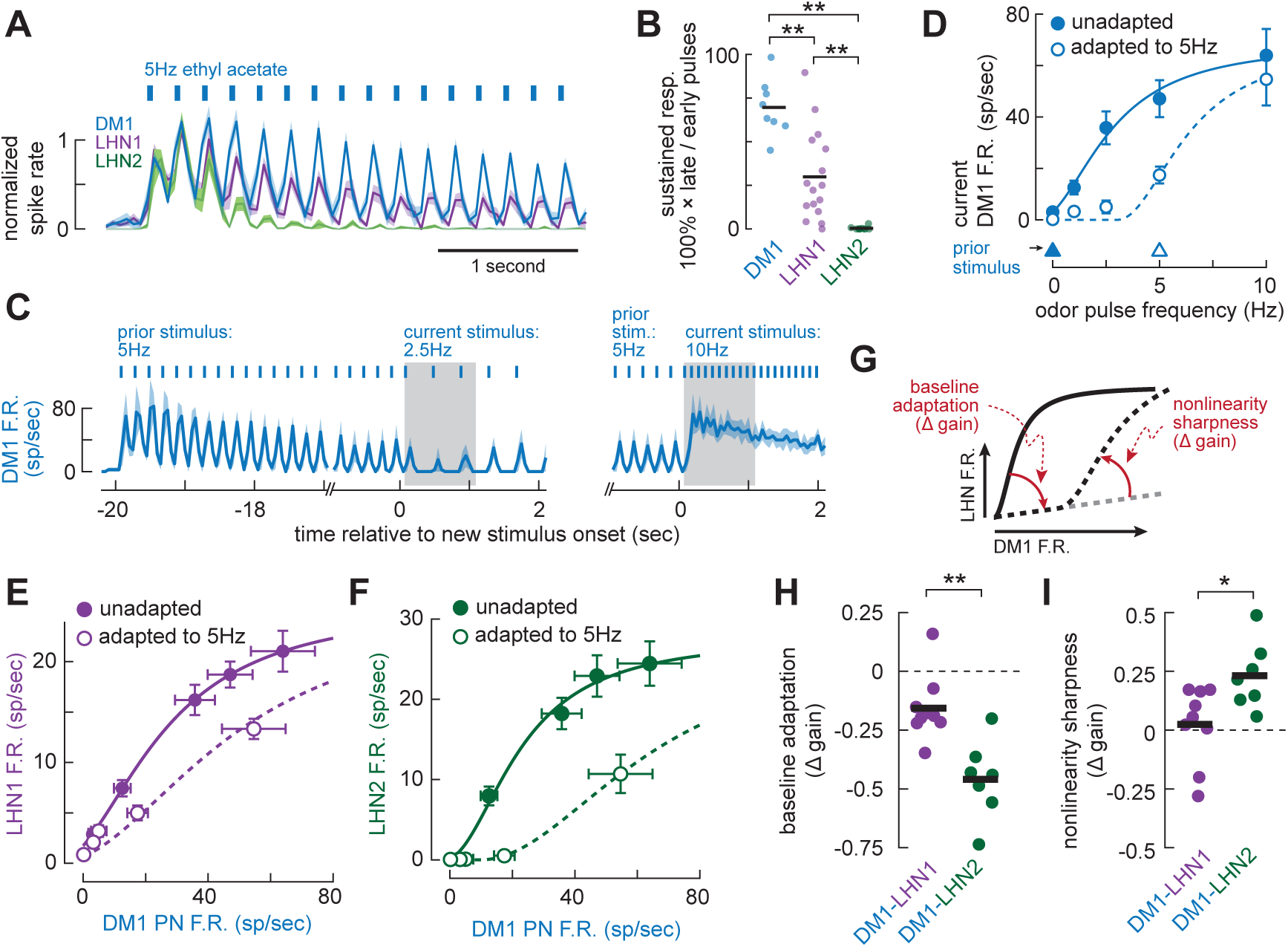
LHN adaptation further specializes upon pre-adapted PN input. **(A)** Mean (± s.e.m.) normalized spike rates of DM1 (n = 8), LHN1 (n = 16), and LHN2 (n = 10), in response to ethyl acetate pulsed at 5Hz. Traces are normalized by the average of the peak spike rates in response to the first two odor pulses. **(B)** Comparison of sustained responsiveness (mean spike rate for odor pulses 19-30 as a percentage of the mean spike rate for odor pulses 1-2) for DM1, LHN1 and LHN2. T tests: ** p < 0.001. **(C)** Mean (± s.e.m.) spike rates in DM1 (n = 4) in response to ethyl acetate pulsed at 5Hz for 20 seconds prior to switching to either 2.5Hz (left) or 10Hz (right). **(D)** Ethyl acetate tuning curves for DM1 spike rates (mean ± s.e.m., n = 4-8) without adaptation (solid circles) and adapted to 5Hz (open circles). Curves were fit as in Figure 1G,H. Prior pulse frequencies are plotted in triangles at bottom. Adapted responses were best fit with a pure subtractive shift, but responses to low stimulus frequencies were notably underpredicted by this model. **(E)** Input-output functions relating DM1 spike rates (mean ± s.e.m., n = 4-8) to LHN1 spike rates (mean ± s.e.m., n = 9-16), without adaptation (solid circles), and after adaptation to 5Hz ethyl acetate (open circles). **(F)** Same as (D) but for LHN2 (n = 6-16). **(G)** Schematic of parameters quantified from input-output functions. Baseline adaptation was measured as the difference in slope between unadapted and 5Hz adapted input-output functions (up to DM1 spike rates of 17Hz). The sharpness of the adapted nonlinearity was measured as the difference in slope between the adapted low-end range (i.e., up to 17Hz DM1 spike rates) and the adapted high-end range. **(H)** Comparison of baseline adaptation for DM1-LHN1 and DM1-LHN2 input-output functions. LHN2 adapts more than LHN1 (** t-test, p = 0.0014). **(I)** Comparison of nonlinearity sharpness for DM1-LHN1 and DM1-LHN2 input-output functions. LHN2 has a sharper nonlinearity than LHN1 (* t-test, p = 0.019).

To isolate the specialization applied by each LHN type to DM1 input, we needed to directly compare adapted responses in each cell type across a range of stimuli. Thus, we measured DM1 spike rates to the same odor pulse frequency increases and decreases that we used to characterize LHNs. DM1 spikes faithfully tracked each odor pulse during increases and decreases in stimulus frequency (**Figure 2C**). After adapting to 5Hz prior stimulation, average DM1 spike rates were broadly reduced, except when odor frequency increased to 10Hz (**Figure 2D**). Viewed as a function of DM1 spike rates, LHN1 itself imposed a modest reduction in gain (**Figure 2E**). In contrast, LHN2 imposed stronger adaptation that included a substantial rightward shift, leading to a strong expansive nonlinearity (**Figure 2F**).

For low DM1 rates (up to ∼20Hz), adaptation was stronger for LHN2 than for LHN1, with LHN2 adapting almost completely (**Figure 2G,H**). However, LHN2’s adapted response function became highly nonlinear at higher DM1 rates, while LHN1 remained nearly linear (**Figure 2G,I**). Similar distinctions in adaptation were also seen for inputs from DM4 (**Figure S3A-G**). LHN2 was also distinct in this circuit because it evoked almost no spontaneous spikes (**Figure S3H**). It thus has a propensity to become completely silent, suppressing both background activity and predictable odor-evoked activity.

### Adaptation in LHN2 is more input specific than adaptation in LHN1

To further constrain the loci of LHN adaptation, we performed cross-adaptation experiments. If input from one PN reduces LHN responses to subsequent input from a different PN, then adaptation is likely implemented by a postsynaptic cellular mechanism. If not, then adaptation is likely implemented by an input-specific (e.g., synaptic) mechanism.

To carry out these experiments, we took advantage of the fact that both LHN1 and LHN2 receive input from both DM1 and DM4 PNs (**Figure S1A,B,S4A**). As previously noted, DM4 is privately activated by low concentrations of methyl acetate (**Figure S1E-G**). We verified that ethyl acetate did not cross-adapt methyl acetate responses in DM4 PNs (**Figure S4B**). Interestingly, ethyl acetate led to a significantly larger reduction in subsequent responses to methyl acetate in LHN1 than in LHN2 (**Figure S4C,D**). In other words, cross adaptation was smaller – and therefore more input-specific – for LHN2 than for LHN1.

### PN-LHN synapses exhibit target-cell-specific short-term plasticity

Differences in the input-specificity of adaptation led us to hypothesize that synapses from DM1 onto LHN1 operate differently from synapses onto LHN2. Specifically, short-term plasticity might differ for different target neurons^5-9^. We thus used paired recordings to compare the physiology of DM1 synapses onto each LHN type.

To mimic odor-evoked PN activity, we delivered 100msec pulses of 100-300pA depolarizing current at 5Hz into DM1, which evoked about 6-8 spikes per pulse (**Figure 3A,B**, bottom traces, **STAR Methods**). Simultaneous recordings from LHN1 revealed large responses to each pulse that declined minimally over repeated pulses. **Figure 3A**, top traces). This is consistent with previous reports of large and reliable unitary excitatory postsynaptic potentials (uEPSPs) at this connection^13,34^. Simultaneous recordings from DM1 and LHN2 revealed moderate responses to the first few pulses which then faded away over subsequent pulses (**Figure 3B**, top traces). Notably, every DM1 spike evoked a clear uEPSP in LHN1 (**Figure 3A**, right) whereas multiple DM1 spikes were required before measurable uEPSPs were evoked in LHN2 (**Figure 3B**, right).

**Figure 3:**
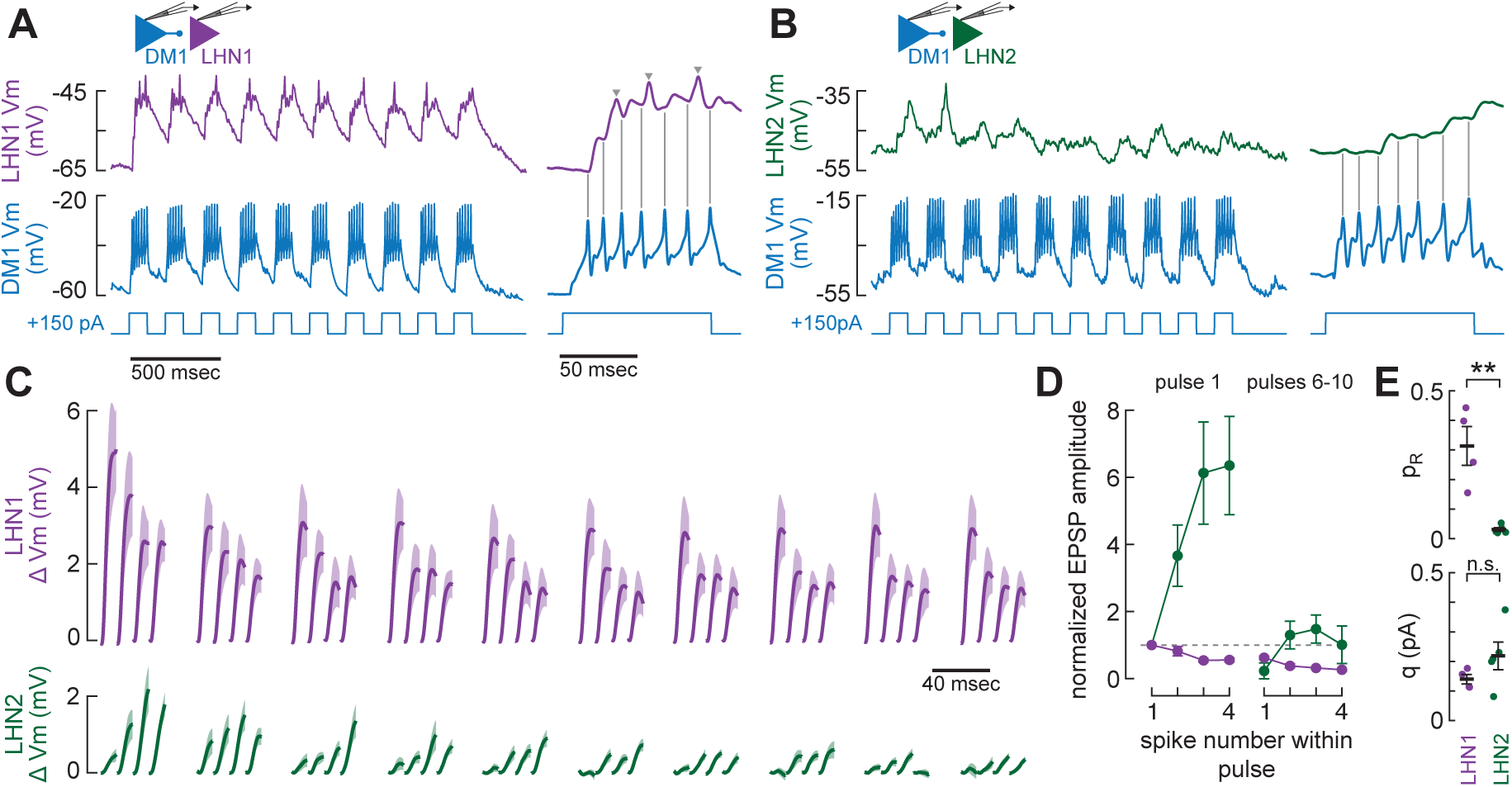
PN-LHN synapses exhibit target-cell-specific short-term plasticity. **(A)** Example paired recording from DM1 and LHN1 showing applied current-evoked DM1 spikes (to mimic 5Hz odor pulse responses) and corresponding postsynaptic responses in LHN1. Right: close up of the response to the first current pulse. Gray arrowheads denote LHN action potentials. **(B)** Same as (A), but for DM1 and LHN2. **(C)** Mean (± s.e.m.) uEPSP waveforms measured in LHN1 (n = 4 pairs) or LHN2 (n = 6 pairs) for the first four DM1 spikes of each current pulse. Each uEPSP waveform is displayed on the same timescale, but each uEPSP is spaced uniformly for clarity. The gap between pulses is not to scale. **(D)** Normalized uEPSP amplitude for DM1-LHN1 (purple) and DM1-LHN2 (green) synapses for the first four spikes of the first pulse, and the first four spikes averaged over the last 5 pulses. **(E)** Initial probability of release (top) and quantal content (bottom) for synapses from DM1 onto LHN1 and LHN2 estimated from paired recordings using quantal theory and inference of the number of release sites from the hemibrain connectome (STAR Methods). t test: ** p < 0.005.

To study the short-term plasticity of each synapse type, we compared the average uEPSP waveforms corresponding to the first four DM1 spikes of each current pulse. uEPSPs in LHN1 were initially very large (∼5mV) but then depressed by about 50% over subsequent spikes within each pulse (**Figure 3C**, top). The 100msec pause between current pulses was insufficient for complete recovery, so the first uEPSP in subsequent pulses was smaller than in the first pulse. Even at the end of 10 pulses, this synapse was still capable of strong depolarization because the initial uEPSP amplitude was so large (**Figure 3C,D**). Conversely, uEPSPs in LHN2 were initially very small (<0.5mV) but facilitated rapidly over subsequent spikes within the first current injection pulse (**Figure 3C**, bottom). During later pulses, the intra-pulse facilitation waned (and ultimately disappeared), leading to uEPSP amplitudes similar to the first uEPSP in the first pulse (**Figure 3C,D**). Quantal analysis (**STAR Methods**) predicted substantially higher initial release probability for synapses onto LHN1 than onto LHN2, with no change in quantal size (**Figure 3E**).

In short, DM1 output synapses have different dynamics for different target LHNs. The DM1-LHN1 synapse is initially strong but exhibits short-term depression. The DM1-LHN2 synapse is initially weak but exhibits short-term facilitation that eventually subsides with repeated DM1 spikes. The net effects of these short-term plasticity differences are well aligned with the properties of odor-evoked dynamics observed in each LHN type.

### PN-LHN synapses exhibit target-cell-specific sensitivity to presynaptic calcium buffering

Presynaptic mechanisms typically implement short-term plasticity^35^, with facilitation classically arising from accumulation of presynaptic calcium^36^. Accordingly, presynaptic dialysis with the slow calcium buffer EGTA selectively blocks short-term facilitation while leaving short-term depression unaffected^6^. EGTA can thus reveal dependence of each type of DM1 output synapse on slow accumulation of presynaptic calcium.

We repeated the same experiments as above, except we included 1mM EGTA in the DM1 patch pipette. (Note that this is frequently included in “standard” internal solution recipes for *Drosophila*, **STAR Methods**.) After allowing ∼15 minutes for EGTA to dialyze into DM1 axon terminals, we observed no substantial changes in LHN1 responses to DM1 spikes evoked by current injection (**Figure 4A,B**). Similarly, no changes over the same time window were observed when EGTA was omitted from the pipette (**Figure 4C**).

**Figure 4:**
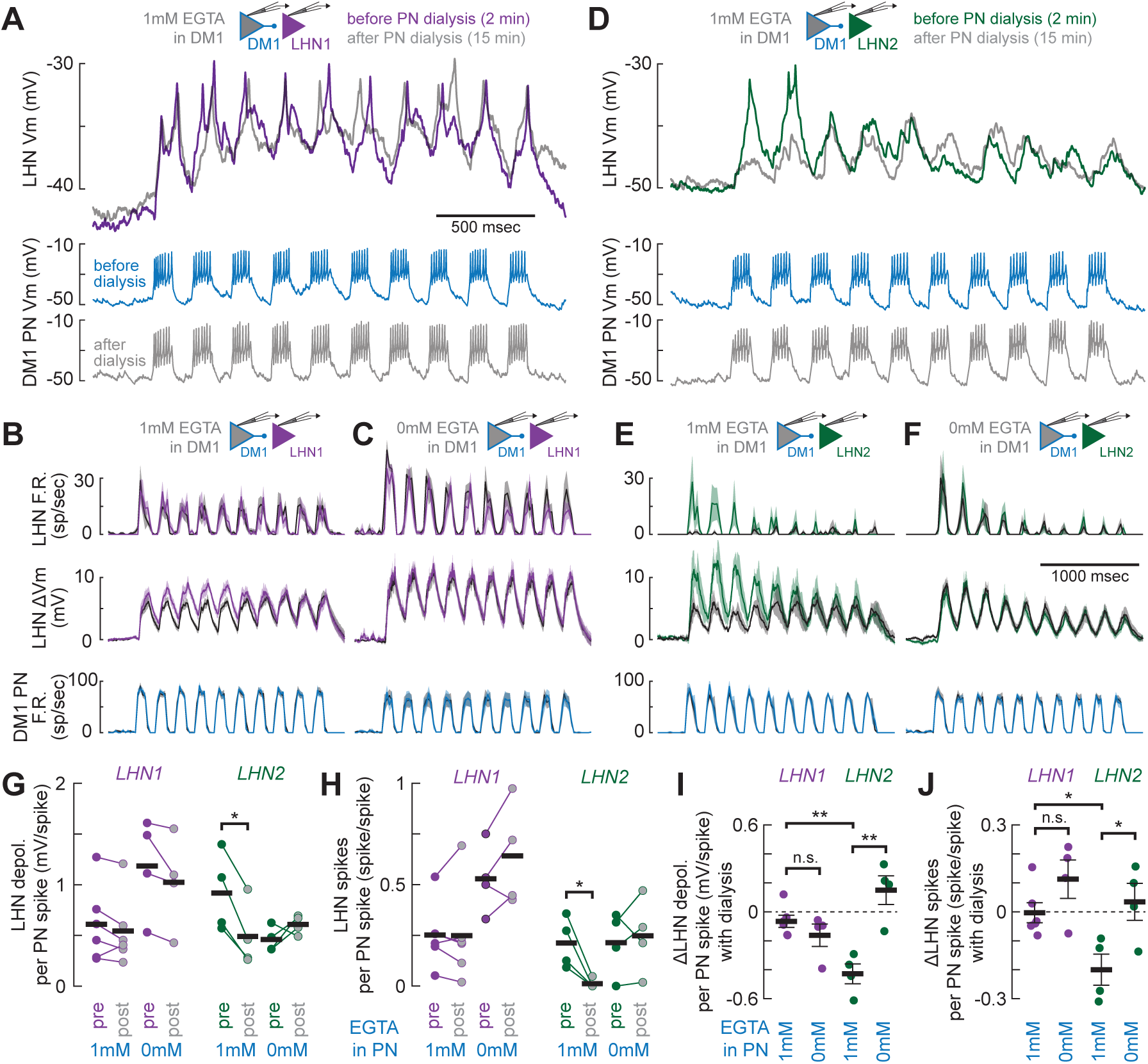
PN-LHN synapses exhibit differential sensitivity to presynaptic calcium buffering. **(A)** Top: individual example traces of LHN1 activity before (purple) and after (gray) PN dialysis with 1mM EGTA. Bottom: example traces of PN activity (simultaneously recorded with the LHN1 traces at top) before (blue) and after (gray) dialysis (evoked via current injection as in Figure 1A,B). **(B)** Bottom: Mean (± s.e.m.) DM1 spike rates evoked by applied current before (blue) and after (gray) dialysis with 1mM EGTA. Middle: mean (± s.e.m.) LHN1 membrane potential before (purple) and after (gray) PN dialysis. Top: mean (± s.e.m.) LHN1 spike rate before (purple) and after (gray) PN dialysis (n = 6 pairs). **(C)** Same as (B) but without EGTA in the PN pipette (n = 4 pairs). **(D)** Same as (A), but for LHN2. **(E)** Same as (B), but for LHN2 (n = 3 pairs). **(F)** Same as (C), but for LHN2 (n = 4 pairs). **(G)** LHN1 depolarization per PN spike (during the first applied current pulse) did not change following PN dialysis with EGTA, but LHN2 depolarization did (paired t-test, p = 0.008). PN dialysis with internal saline lacking EGTA had no significant effect on LHN1 or LHN2 depolarization. **(H)** LHN1 spikes per PN spike (during the first applied current pulse) did not change following PN dialysis with EGTA, but LHN2 spikes did (paired t-test, p = 0.03). PN dialysis with internal saline lacking EGTA had no significant effect on LHN1 or LHN2 spikes per PN spike. **(I)** Dialysis with EGTA reduces the depolarization per PN spike for LHN2 more than without EGTA (t-test, p = 0.003). There is no corresponding difference for LHN1 (t-test, p = 0.27). The EGTA-induced reduction for LHN2 is larger than it is for LHN1 (t-test, p = 0.001). **(J)** Dialysis with EGTA reduces the number of LHN spikes per PN spike for LHN2 more than without EGTA (t-test, p = 0.03). There is no corresponding difference for LHN1 (t-test, p = 0.13). The EGTA-induced reduction for LHN2 is larger

After DM1 dialysis during LHN2 recordings, the initially strong responses during the first two applied current pulses disappeared (**Figure 4D**). EGTA reduced the net LHN2 depolarization and spike rate (**Figure 4E**), but dialysis with internal saline lacking EGTA did not (**Figure 4F-H**). Overall, the net effect of EGTA was significantly greater for LHN2 than for LHN1 (**Figure 4I,J**). Interestingly, EGTA in DM1 also made LHN2 response dynamics resemble LHN1 response dynamics. This suggests that synapses onto LHN2 have an additional pathway of release – not present in synapses onto LHN1 – which may be entirely responsible for transient facilitation.

These experiments identify differences in calcium dynamics at different presynaptic terminals of the same PN axon. However, these terminals are not physically separated: single axon branches form synapses onto both LHN types (**Figure S5**). This means that facilitation and depression are likely determined by the postsynaptic neuron (rather than axon branch specific regulation or cell-extrinsic spatial gradients) but are implemented by the presynaptic neuron.

### Unc13B-mediated synaptic transmission enables temporal prediction error coding in LHN2

The essence of the prediction error code in LHN2 comes from its sharp nonlinearity after adaptation, briefly boosting responses to sudden increases in PN spike rate (**Figure 2**). The transient facilitation (**Figure 3**) and EGTA-dependent boost of synaptic transmission (**Figure 4**) at the onset of DM1 spiking suggests that PN-LHN synapses directly implement this nonlinearity. We thus sought to identify its molecular basis.

EGTA-sensitive synaptic facilitation typically requires long coupling distances between presynaptic calcium channels and calcium sensors^6,37-39^. This is because EGTA is too slow to affect closely coupled release sites, where calcium does not need much time to reach the sensor^37^. Coupling distances are mediated by presynaptic Unc13 proteins, which also impact short-term plasticity^40^. In *Drosophila*, the Unc13B isoform specifically mediates long coupling distances, participates at facilitating synapses, and is present in the lateral horn^41-43^. Therefore, we hypothesized that Unc13B underlies LHN2’s nonlinear response function by transiently elevating synaptic output only when DM1 activity increases (**Figure 5A**).

**Figure 5:**
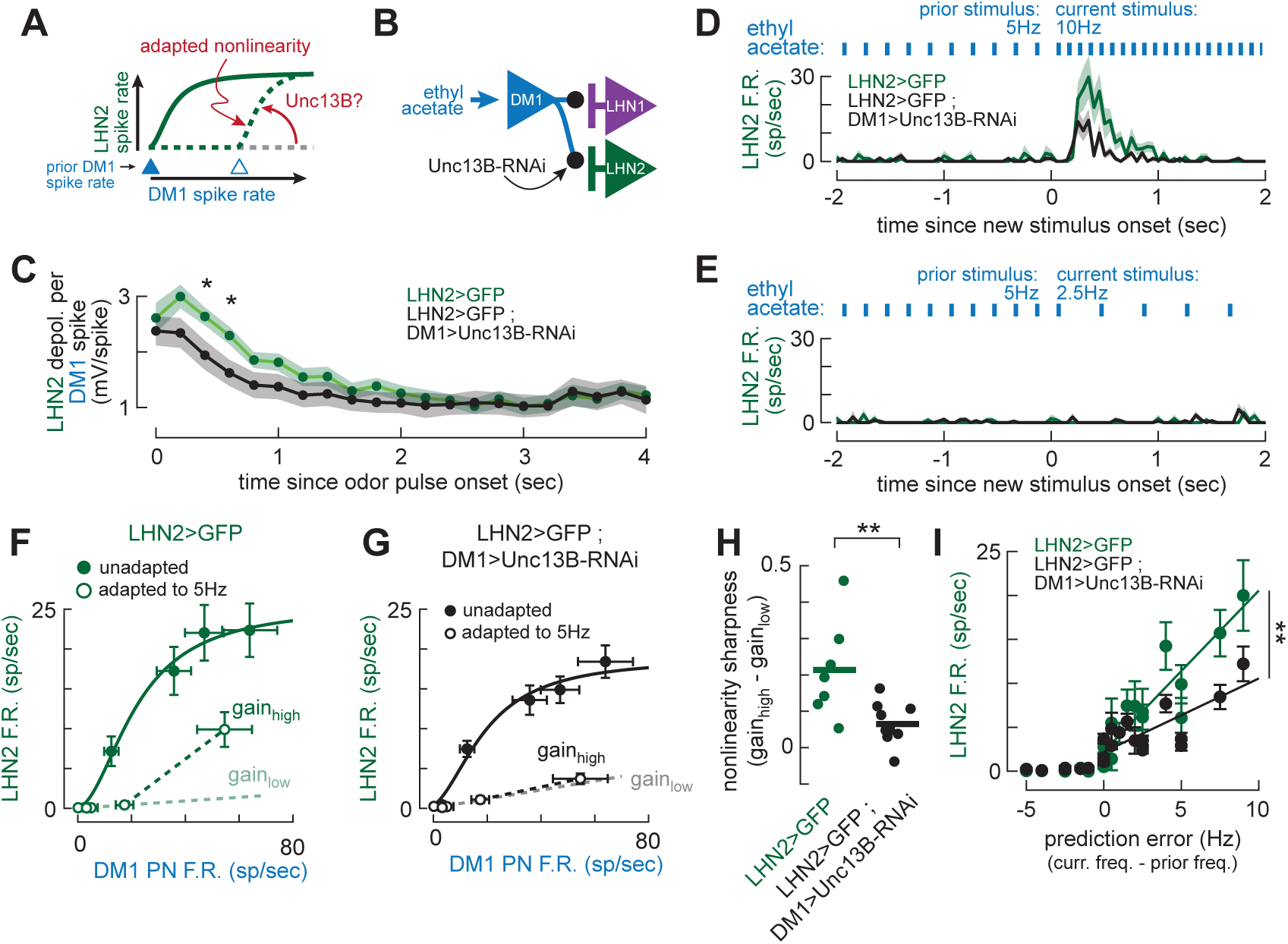
Unc13B-mediated synaptic transmission enables temporal prediction error coding in LHN2. **(A)** Schematic of hypothesis tested here, that Unc13B in PN axon terminals implements the sharp adapted nonlinearity in LHN2. **(B)** Schematic of experimental manipulation knocking down Unc13B expression in PN axon terminals. **(C)** Mean (shading denotes s.e.m.) net LHN2 depolarization per spike per 5Hz odor pulse. Each point denotes one odor pulse in control conditions (green, n = 15) and with DM1 Unc13B knockdown (black, n = 18). T-tests, * p < 0.05. **(D)** Mean (± s.e.m.) PSTHs of LHN2 responses to ethyl acetate pulse frequency increasing from 5Hz to 10Hz. In control conditions (green, n = 7), transient LHN2 activity is larger than with DM1 Unc13B knockdown (black, n = 10). Only the last 2 seconds of the 20-second prior 5Hz stimulus are shown. **(E)** As in (D) but for ethyl acetate pulse frequency decreasing from 5Hz to 2.5Hz. **(F)** Input-output functions relating mean (± s.e.m.) DM1 spike rates (n = 4-8) to mean (± s.e.m.) LHN2 spike rates, without adaptation (solid circles, n = 10-12) and after adaptation to 5Hz ethyl acetate (open circles, n = 7). The adapted responses were fit using separate linear models for odor pulse rates above the adapting frequency (gainhigh, solid line) and below the adapting frequency (gainlow, dashed line). Data are repeated from Figure 2E for ease of comparison to (G). **(G)** Input-output functions relating mean (± s.e.m.) DM1 spikes rates to mean (± s.e.m.) LHN2 spike rates, as in (F), but with Unc13B knockdown in DM1 (n = 9 LHN2 recordings). **(H)** Unc13B knockdown reduces the nonlinearity sharpness (difference between gainhigh and gainlow). t-test, p = 0.006). **(I)** Mean (± s.e.m.) LHN2 spike rates plotted as a function of prediction error (difference between current stimulus frequency and prior frequency), for control conditions and with Unc13B knockdown in DM1. Unc13B knockdown significantly reduces the slope for positive prediction errors (ANCOVA interaction, p = 0.0002).

To test this hypothesis, we reduced Unc13B expression in DM1 using RNAi (**Figure 5B**; **STAR Methods**). This manipulation reduced odor-evoked initial transient depolarizations in LHN2 (**Figure 5C**), but not LHN1 (**Figure S6**), mimicking EGTA dialysis. It also did not affect methyl acetate responses in LHN2, or intrinsic excitability in LHN2, indicating that this manipulation is indeed selective for DM1 inputs (**Figure S6**). Unc13B-RNAi substantially reduced LHN2 spike responses to the increase in odor pulse rate from 5Hz to 10Hz (**Figure 5D**). Spike responses to steady state odor pulses or decreases in odor pulse rate remained nearly absent (**Figure 5E**). Consequently, the sharp nonlinearity of the adapted DM1-LHN2 input-output function was abolished with knockdown of Unc13B (**Figure 5F-H**).

Disrupting this nonlinearity should also impair the fidelity of LHN2’s prediction error code. Indeed, LHN2 spike rates grew more gradually with prediction error when Unc13B expression was reduced (**Figure 5I**). In other words, LHN2 used less of its dynamic range to encode the same prediction error values. Collectively, these data show that Unc13B enables a transient component of synaptic transmission during unexpected increases in PN activity, which contributes to the adaptive nonlinearity necessary for encoding positive prediction error.

### The Na+/K+ ATPase enables temporal contrast coding in LHN1

Temporal contrast coding in LHN1 arises from gain control that depends on the strength of prior stimulation (**Figure 1G,I**). Because LHN1 cross-adapts across different PN inputs (**Figure S4**), LHN1 likely controls its gain intrinsically. However, this requires maintaining a running average of recent activity to use for attenuating responses to future synaptic inputs. How does LHN1 achieve this?

A first clue to the identity of this mechanism was that LHN1 tends to increasingly hyperpolarize after each odor pulse (when spaced far enough apart; **Figure 6A**, top). Hyperpolarization was also evident after pulses of current injection, suggesting a cellular origin because LHN1 has almost no feedback connections (**Figure S7**). These membrane potential dynamics resembled the electrogenic properties of the Na+/K+ ATPase (hereafter referred to simply as the sodium pump) which induces an outward current as it pumps sodium out of the cell^44-46^. High levels of activity can elevate intracellular sodium concentration enough to require seconds to tens of seconds of pump activity to restore it to baseline^47^. Sodium thus stores a cellular memory of recent activity, which is “recalled” by the action of the sodium pump^45^. We thus hypothesized that this implements gain control in LHN1 (**Figure 6B**).

**Figure 6:**
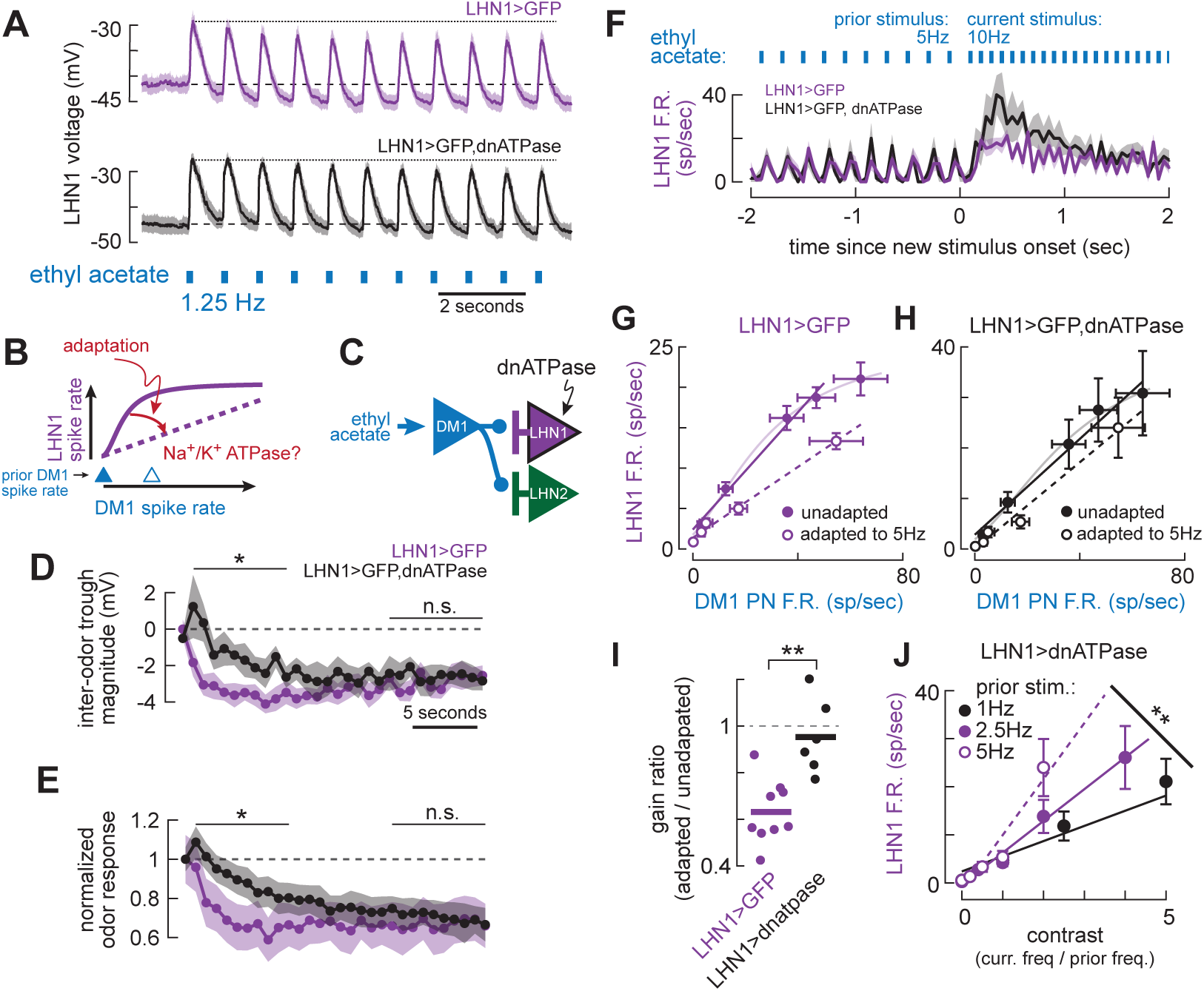
The Na+/K+ ATPase enables temporal contrast coding in LHN1. **(A)** Mean (± s.e.m.) voltage traces in LHN1 in response to 1.25Hz ethyl acetate pulses. Top, control (n = 11). Bottom: LHN1>dnATPase (n = 7). **(B)** Schematic of hypothesis tested here, that the Na+/K+ ATPase underlies temporal gain control and contrast coding in LHN1. **(C)** Schematic of experimental manipulation. **(D)** Mean (± s.e.m.) inter-odor trough voltage (relative to baseline) after each odor pulse for the traces in (A), for control (purple) and LHN1>dnATPase (black). Each point denotes the trough following each odor pulse. Mean voltage in pulses 2-10 is higher for LHN>dnATPase than for control (t-test, p = 0.016), but not in pulses 21-30. **(E)** As in (D), except for the average depolarization for each odor pulse (normalized to the amplitude for the first pulse). The mean response in pulses 2-10 is higher for LHN>dnATPase than for control (t-test, p = 2.5 × 10^-4^), but not in pulses 21-30. **(F)** Mean (± s.e.m.) PSTHs of LHN1 responses to ethyl acetate pulse frequency increasing from 5Hz to 10Hz. In control conditions (purple, n = 8), transient LHN1 activity is smaller than with expression of dnATPase (black, n= 6). Only the last 2 seconds of the 20-second prior 5Hz stimulus are shown. **(G)** Input-output functions relating mean (± s.e.m.) DM1 spike rates (n = 4-8) to mean (± s.e.m.) LHN1 spike rates, without adaptation (solid circles, n = 5-16) and adapted to 5Hz ethyl acetate (open circles, n = 9). Error bars denote s.e.m. The linear regime of the unadapted (solid purple line) and adapted (dashed line) responses were fit with linear models. The full nonlinear model is shown in the pale purple line for comparison. Data are repeated from Figure 2E for ease of comparison to (H). **(H)** Input-output functions relating DM1 spikes rates to LHN1 spike rates, as in (F), but with dnATPase expression in LHN1 (n = 6 LHN recordings). **(I)** dnATPase expression in LHN1 reduces the gain ratio (adapted / unadapted) of the DM1-LHN1 input-output function (** t-test, p = 0.001). **(J)** Mean (± s.e.m.) LHN1 spike rates as a function of temporal contrast (current pulse frequency divided by prior frequency). Linear fits for each prior frequency have different slopes (** ANCOVA interaction, p = 0.0004), unlike the case for control LHN1 recordings, where slopes are indistinguishable (Figure 1I).

To test this hypothesis, we genetically perturbed the sodium pump by overexpressing a dominant negative version with a mutant alpha subunit (“dnATPase”) selectively in LHN1^48^ (**Figure 6C**). This manipulation reduces the number of functional sodium pump molecules because they get out-competed by the dominant negative version^45,48^. Expression of dnATPase caused inter-odor hyperpolarizations to grow more slowly over time (**Figure 6A,D**), allowing responses to the first few odor pulses to stay larger for longer (**Figure 6A,E**). This suggests that the outward current from the sodium pump contributes to adaptation, in part, by narrowing the temporal integration window for incoming PN spikes.

With fewer functional pump molecules, LHN1 should be more responsive to increases in odor pulse frequency. This is because the remaining pump current will be slower to adjust to the new stimulation level, so an increase in PN spike rate will encounter a temporal integration window matched to the lower level of activity. In agreement with this, expression of dnATPase transiently elevated LHN1 spike rates when the odor pulse rate increased from 5Hz to 10Hz (**Figure 6F**). More broadly, it also eliminated the gain reduction of the DM1-LHN1 input-output function after adaptation to 5Hz odor pulses (**Figure 6G-I**).

Since contrast coding depends on gain modulation in LHN1, it should also be disrupted by expression of dnATPase. Accordingly, LHN1 spike rates no longer formed a single function of temporal contrast (**Figure 6J**; compare to Figure 1I). In short, by taking advantage of the slow and activity-dependent dynamics of intracellular sodium, the sodium pump counteracts excitation to reduce gain on slow timescales. In LHN1, this contributes to the implementation of a temporal contrast code.

### LHN1 and LHN2 contribute distinct dynamics to behavioral attraction to food odor

LHN1 and LHN2 both contribute to innate olfactory attraction^21,27^. The distinct dynamics of their physiological responses to odors suggest that their behavioral contributions may also be dynamically distinct. Because these are just two of the hundreds of LHN types, and individual LHN types typically make relatively small contributions^21^, we focused our behavioral experiments on the simple difference in transience between LHN1 and LHN2. Accordingly, we hypothesized that LHN1 has a sustained effect on odor attraction, while LHN2 has a transient effect.

We tethered flies to walk on a spherical treadmill while delivering food odors with temporal precision to the front of the fly. In this configuration, behavioral attraction is evident as a relative increase in walking speed. We then used selective split-Gal4 lines^21^ to silence synaptic output of either LHN1 or LHN2 with the temperature sensitive dynamin *shibire*^49^ (**STAR Methods**). Because robust odor-evoked navigation usually requires activity in many PNs, we used apple cider vinegar as a stimulus, which indeed evoked large behavioral responses in control flies. Importantly, the physiological differences in transience between LHNs occurred for all stimuli tested, including those that activated multiple PNs (**Figure S2**).

Silencing LHN1 reduced the relative increases in walking speed in response to both a single long pulse of high concentration vinegar and intermittent short pulses of low concentration vinegar (**Figure 7A**). Significant reductions were observed both early in the stimulus and late in the stimulus (**Figure 7B**). Thus, LHN1 made sustained contributions to odor attraction.

**Figure 7:**
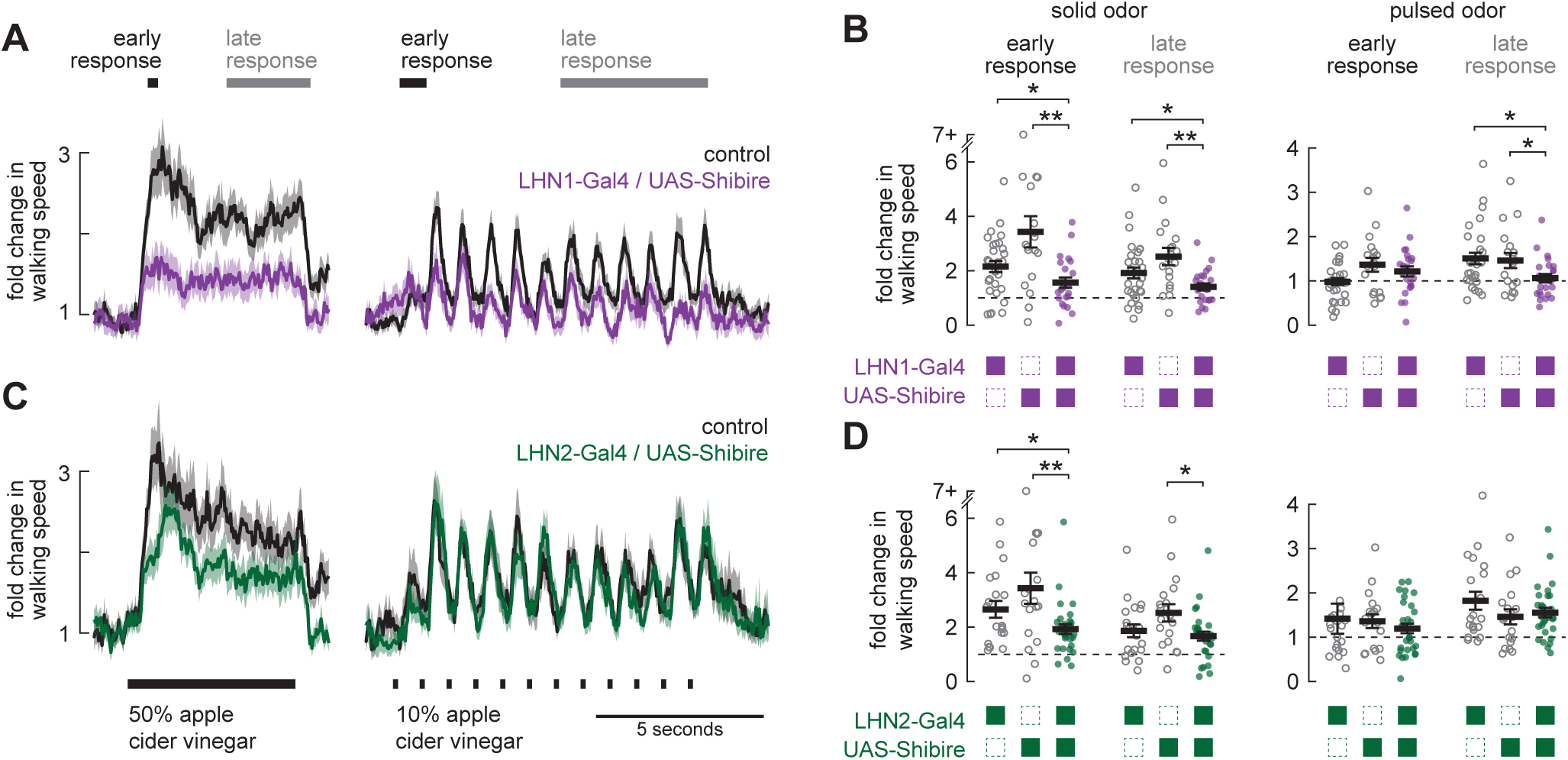
LHN1 and LHN2 contribute distinct dynamics to behavioral attraction to food odor. **(A)** Mean (± s.e.m.) fold change in walking speed during presentation of solid 50% apple cider vinegar (left) or pulsed 10% apple cider vinegar (right). Early and late response windows denote periods used for analysis in panel C. Black traces are the pooled Gal4 and UAS controls, purple traces are with shibire expressed in LHN1. **(B)** Quantification of mean response in early and late time windows (as denoted in panel A), for each fly of each control genotype and for the genotype expressing shibire in LHN1. Solid black lines and error bars deonte mean and s.e.m. across flies. **(C)** Same as (A) but with shibire expressed in LHN2. **(D)** Same as (B) but for shibire expressed in LHN2 t-tests: * p < 0.05; ** p < 0.005.

Silencing LHN2 only reduced the relative increases in walking speed during a narrow time window at the onset of the long pulse of high concentration vinegar (**Figure 7C**). Behavior was not significantly affected during any other component of either stimulus (**Figure 7D**). LHN2 therefore makes transient contributions to odor attraction, but only for strong enough stimuli.

These experiments illustrate that the behavioral roles of each LHN type correspond to their neural coding properties. Thus, the divergent processing that begins with subcellular synaptic specializations in PNs creates parallel pathways that can independently control different dynamic components of behavior.

## DISCUSSION

By its nature, sensory processing in the brain discards information. Since the same information may be relevant for some purposes and not for others, a widespread strategy is to split sensory processing into parallel pathways so that each pathway can specialize^50^. Our results demonstrate that subcellular presynaptic specializations originate diverse temporal coding in parallel channels. This allows predictable stimuli to be discarded from one pathway but preserved in another, allowing for greater behavioral flexibility.

### Dynamics of higher-order olfactory coding

The receptive field of a third-order olfactory neuron is essentially a weighted sampling of glomeruli, including both positive and negative weights^13^. Different sampling yields neurons tuned to different regions of olfactory space^14,15^. Here, we show that this principle extends to the time domain. Glomerular weights can rapidly change based on recent odor exposure, yielding some neurons that faithfully track all temporal patterns (LHN1), and other neurons that only detect increases in odor intensity (LHN2).

The structure of glomerular sampling by LHNs is nonrandom, and many of the same combinations of glomeruli are co-sampled by different LHN types^12,14^. Although co-sampling is not typically identical, our results provide an additional explanation for this partial redundancy: different dynamics of the same odor patterns are encoded in different LHNs. In this regard, it is interesting that adaptation is highly stereotyped for each LHN type, in parallel with the stereotypy of glomerular sampling. This suggests that synaptic connectivity and synaptic dynamics are both genetically specified in this circuit.

Our data show that the sustained and transient LHN representations separately control corresponding sustained and transient components of a “single” behavior (walking speed) in response to odor pulses. Notably, silencing LHN1 had overall a larger impact than silencing LHN2, which might simply reflect the fact that our paradigm did not require any intrinsically transient behavior. It will thus be interesting to learn whether LHN2 contributes more directly to intrinsically transient behaviors, such as turning upwind after first making contact with a food odor^51,52^.

Because most LHNs are at least several synapses away from motor control centers, other functions of temporal coding diversity may be less visible to direct behavioral measurement. Interestingly, a 10-fold expansion of cell types occurs in the lateral horn, where ∼50 uniglomerular PN types contact ∼500 LHN types^12^. One possibility is that the lateral horn as a whole forms a “chemotemporal” basis set, consisting of various combinations of molecular features and temporal dynamics, which might then combine to link specific combinations of chemical dynamics to specific instinctive behavioral programs. In support of this idea, broad chemotemporal diversity has been reported across LHNs in both fly^15^ and locust^53^.

### Target-cell specific short-term plasticity originates divergent dynamics

A central finding of our study is that temporal diversity in LHNs arises largely from target-neuron specific short-term plasticity in upstream PNs. Similar to other reports of this phenomenon^6^, we found that subcellular differences in presynaptic calcium dynamics generate this specificity, because PN dialysis with EGTA only disrupted synapses onto LHN2. This means that the same PN spike trains should drive different patterns of neurotransmitter release onto different target neurons. PN output is thus multidimensional: constrained by spike patterns, but not fully defined by them.

We found that PN output synapses that facilitate and then depress underlie transient LHN2 responses to odor. In other circuits, transience derives from purely depressing synapses^54-56^. What advantage does initial facilitation provide? Synapses that depress usually have high initial release probabilities, which strongly transmit low spike rates. Because PNs are spontaneously active^57^, low spike rates may purely be noise. Facilitation blocks this noise, because initial release probabilities can be nearly zero, and spontaneous rates are not high enough to elevate them substantially. Accordingly, LHN2 was almost completely devoid of spontaneous activity, while average DM1, DM4, and LHN1 spontaneous rates were all ∼3Hz. Facilitation thus operates like a noise gate in audio engineering used to suppress weak signals.

During prolonged PN activity, LHN2 spiking fully adapts away, but can rapidly be reawakened by increased PN activity. This, too, benefits from facilitation at this synapse. Depression alone would need to be nearly complete to suppress steady state spiking. If depression were implemented by depletion of vesicle pools, this would severely limit the amount of releasable transmitter available when PN activity increases. However, transient facilitation allows substantial transmitter release in this situation, suggesting that the “activation threshold” of the noise gate is variable and dependent on past activity.

In contrast, we found that sustained LHN1 responses arose from PN output synapses that purely depress. Why does this depression not cause more transience? In large part, transience is avoided because the depression is relatively modest. Even after ∼80 PN spikes within 2 seconds, individual EPSPs are still ∼40% of their initial size, enough to maintain LHN1 depolarization, but perhaps not enough to rapidly initiate it. These dynamics are akin to the throttle needed to accelerate a car up to a stable speed. The specific mechanisms that enable this synaptic depression are not known but may resemble those of the similarly high-fidelity ORN-PN synapse^16,58^.

### Synaptic mechanisms of temporal prediction error coding

Spike activity in LHN2 was consistent with encoding a positive temporal prediction error, in which the predicted signal is subtracted from the actual signal. This requires a rightward shift in an input-output function, creating a sharp (expansive) nonlinearity. Moreover, the predicted signal must be stored (however briefly) for future use^30^. What creates this nonlinearity and where is the predicted signal stored?

Strikingly, we found that the nonlinearity depends almost entirely on Unc13B-mediated transmission. Unc13B enforces long coupling distances between presynaptic calcium channels and calcium sensors^41^, implicating the active zone as the locus of the nonlinearity. The major expansive nonlinearity in the presynapse is the calcium sensor, which scales transmitter release with the third or fourth power of calcium concentration^59^. This nonlinearity is most apparent when calcium is a limiting factor in release, meaning that it is highly relevant for facilitating synapses, which classically rely on calcium accumulation and long coupling distances^35,37^. Because the slow action of EGTA is enough to block DM1-LHN2 facilitation, presynaptic calcium is clearly a limiting factor at this synapse.

We suspect the predicted signal is stored by a process upstream of the calcium sensor, to harness its expansive nonlinearity. A good candidate is the partial inactivation of presynaptic calcium channels, which could reduce presynaptic calcium concentration during steady state activation to levels too low for transmitter release. Subsequent increases in PN activity could then still rapidly raise calcium concentration enough to overcome long coupling distances to evoke strong release^60^. Thus, subtraction could be implemented by engaging different regimes of the calcium sensory nonlinearity.

### Cellular mechanisms of temporal contrast coding

Spike activity in LHN1 was consistent with encoding temporal contrast, meaning that it encodes relative changes in stimulus intensity using linear changes in spike rate. However, since adaptation in LHN1 cross-adapted between PN inputs, synaptic mechanisms (such as depression^54,55,61^) are unlikely to contribute. Instead, we found that temporal contrast coding in LHN1 requires the Na^+^/K^+^ ATPase, which restores ionic balance after strong excitation by pumping out three Na+ ions for every two K+ ions pumped in, causing an outward current.

Intracellular sodium is a convenient way to implement temporal comparisons because it accumulates roughly linearly with neural activity, returns to baseline on timescales of seconds to tens of seconds, and is not highly buffered like calcium^45,47,62^. In this regard, the role of the sodium pump is conceptually similar to Na^+^-dependent K^+^ currents^63^, in that the sodium concentration stores a cellular memory of past activity which is then recalled by hyperpolarizing currents. This might explain why PN-LHN1 synapses do not fully depress despite a presumably large metabolic cost: strong and reliable transmission is necessary to maintain sodium concentrations in an elevated state to accurately reflect the current odor stimulus. It will be useful in the future to make direct measurements of intracellular sodium dynamics to determine how they impact the transformation of synaptic input into spiking output.

### Origination of temporal diversity in central circuits

In other sensory systems, most notably somatosensation, different dynamic representations originate immediately in different peripheral receptor types^64^. Why does the olfactory system postpone temporal diversification? One possibility is the need to detect weak and noisy signals, which benefits from averaging the activity of many ORNs expressing the same receptor^65-68^. Splitting each ORN population into transient and sustained subtypes would yield smaller populations of each, diminishing the benefits of averaging (assuming constraints on the total number of ORNs). Diversifying after averaging thus allows for maximal noise reduction. A similar organization exists in vision, where multiple photoreceptors converge onto bipolar cells, which then generate diverse temporal representations^4^. Originating temporal diversity centrally may thus be a particularly useful strategy when very weak sensory signals are important.

Creating temporal diversity through axonal specializations provides at least two additional advantages. First, it is compact, because it does not require interposed neurons to provide synaptic inhibition, so computation can be performed with fewer neurons. Second, it reduces redundancy as early as possible by suppressing unnecessary synaptic transmission, which reduces metabolic costs^69,70^. Interestingly, specific axonal interactions may also underlie the generation of temporally diverse odor representations in different PNs^71^. This general strategy may thus be widely used to conserve energy and to allow interposed neurons to perform independent functions, such as implementing comparisons across sensory space^3,72,73^.

### Building blocks of dynamic network function

Network connectivity constrains network function, but cannot predict network dynamics^74^. Here, we identify dynamic roles for two proteins – Unc13B and the Na+/K+ ATPase – which diversify the function of a simple divergent network. Expression of Unc13B is not uniform throughout the brain, so its presence at any given synapse may predict dynamics^42^. Mapping Unc13B expression with synaptic resolution could, in principle, constrain the short-term plasticity properties of every synapse in the brain. In contrast, the Na+/K+ ATPase exists in every neuron and its impact on activity likely depends on the dynamics of synaptic inputs. Thus, measuring intracellular sodium dynamics may be as important as calcium dynamics or spike patterns. In the future, it will be important to incorporate all of these dynamic building blocks into connectome-constrained network models to make better predictions of network function and its behavioral consequences.

## ACKNOWLEDGEMENTS

We thank Damon Clark and Evyn Dickinson for help with tethered behavior and analysis, Joel Greenwood and the Kavli Neurotechnology Core at Yale University for technical assistance with behavior and physiology apparatus, Leslie Griffith, Stephan Sigrist, and Damon Clark for sharing fly lines. Michael Higley, Jonathan Demb, Mehmet Fişek, and members of the Jeanne Lab provided helpful comments on the manuscript. This work was supported by NIH grants R01DC018570 and R01NS116584, the Richard and Susan Smith Family Award for Excellence in Biomedical Research, the Klingenstein-Simons Fellowship Award in Neuroscience, and an Innovative Research award from the Kavli Institute for Neuroscience at Yale University to J.M.J. H.S.K. was supported by an NSF Graduate Research Fellowship.

## STAR★METHODS

### KEY RESOURCES TABLE

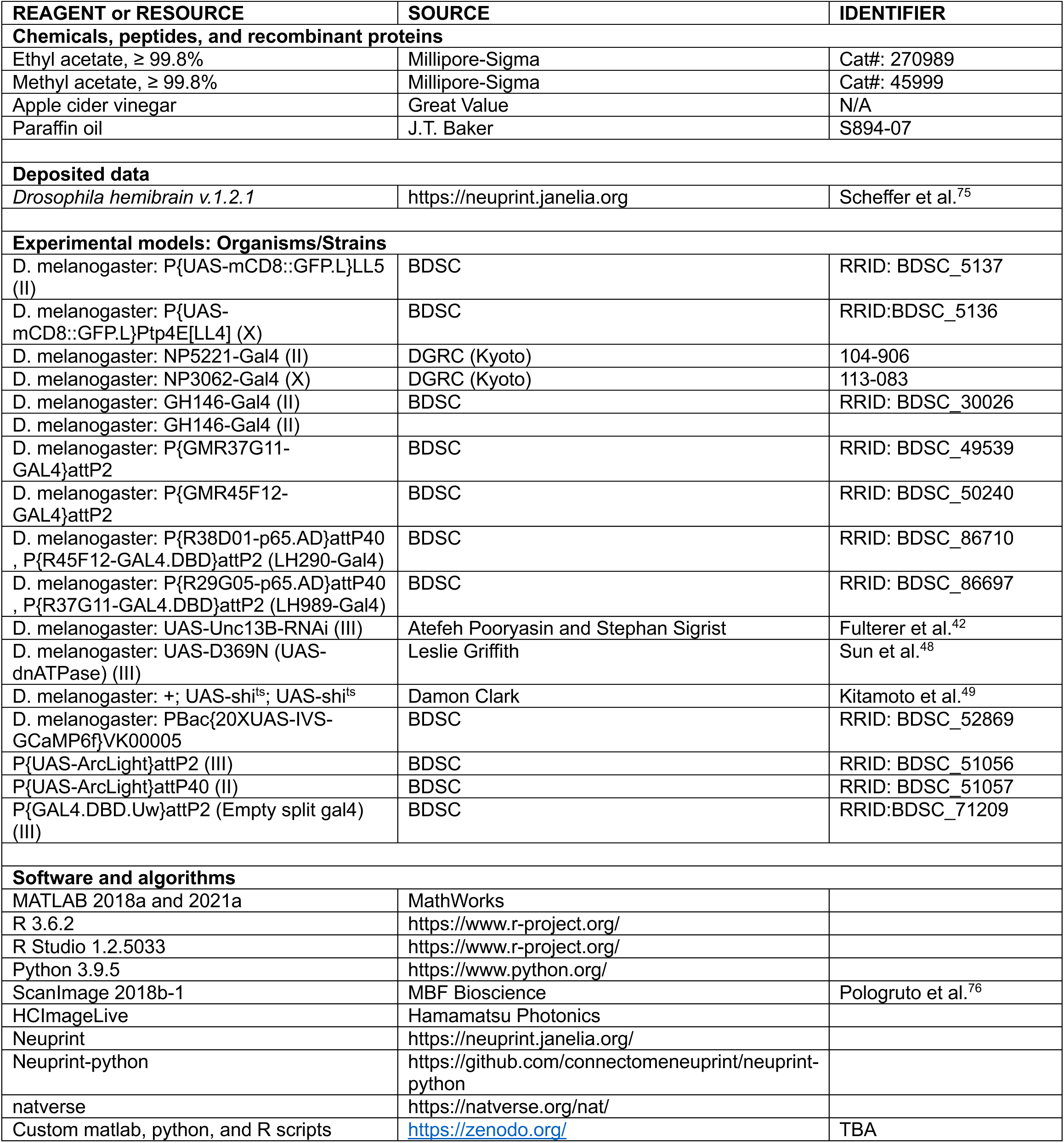

### RESOURCE AVAILABILITY

#### Lead contact

Further information and requests for resources and reagents should be directed to and will be fulfilled by the lead contact, James M. Jeanne.

#### Materials availability

This study did not generate new unique reagents.

#### Data and code availability

Electrophysiology and imaging data will be deposited and publicly available as of the date of publication. Accession numbers are listed in the key resources table.

All original code has been deposited at Zenodo and is publicly available as of the date of publication.

DOIs are listed in the key resources table.

Any additional information required to reanalyze the data reported in this paper is available from the lead contact upon request.

### EXPERIMENTAL MODEL

Flies (*Drosophila melanogaster*) were raised on conventional cornmeal agar medium supplemented with rehydrated potato flakes (Carolina Biological Supply) and yeast under a 12 h light, 12 h dark cycle at 25°C. All electrophysiology experiments were performed *in vivo* on adult female flies either on the day of eclosion (at least 8 hours old) or one day after eclosion. Voltage and calcium imaging experiments were performed on adult female flies 1-4 days after eclosion. Behavior experiments were performed on adult male flies 2-5 days after eclosion, since these flies walked most consistently on the spherical treadmill. None of the neurons in the lateral horn investigated in this study are sexually dimorphic^77,78^.

The following genotypes were used:

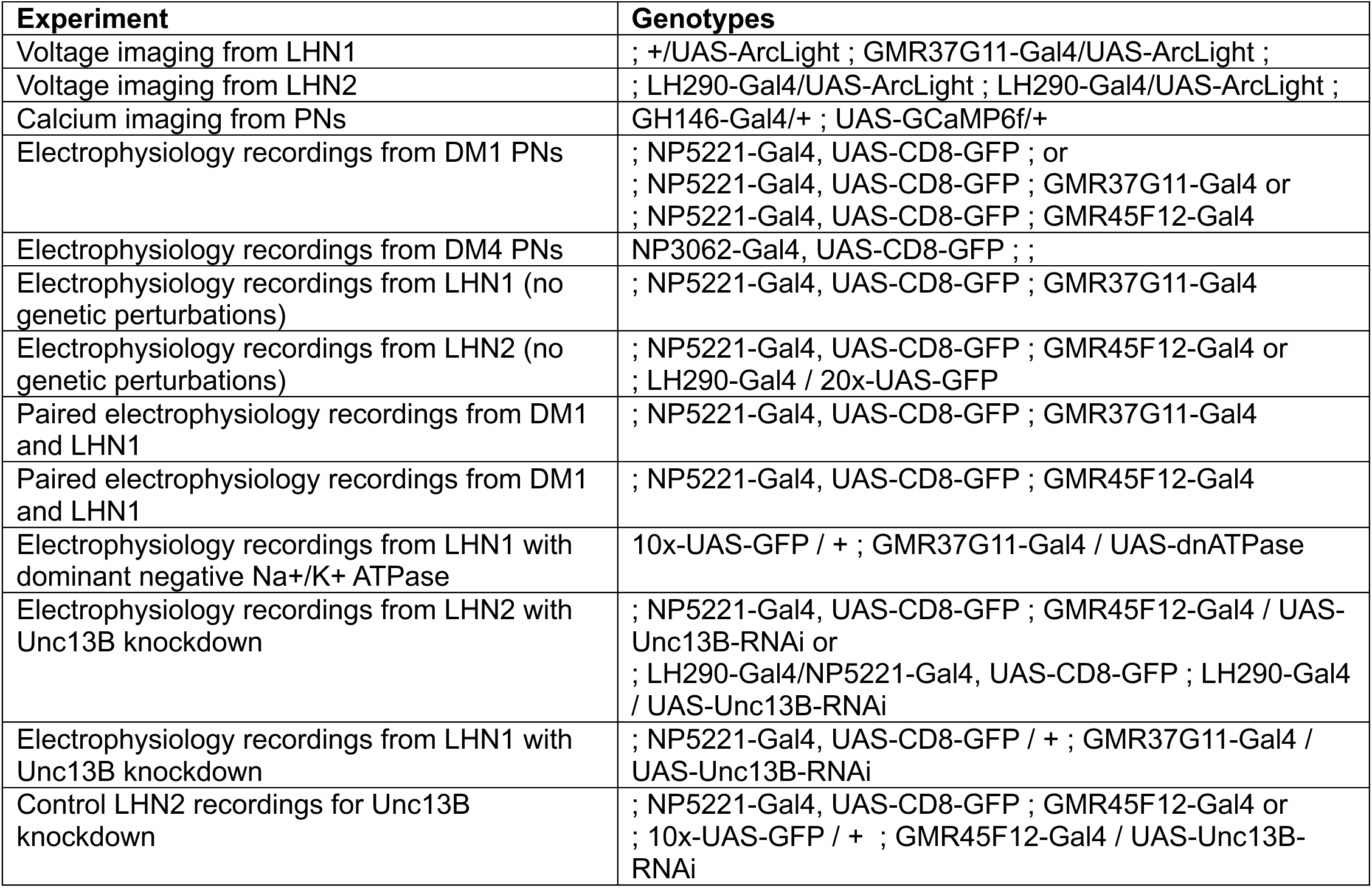
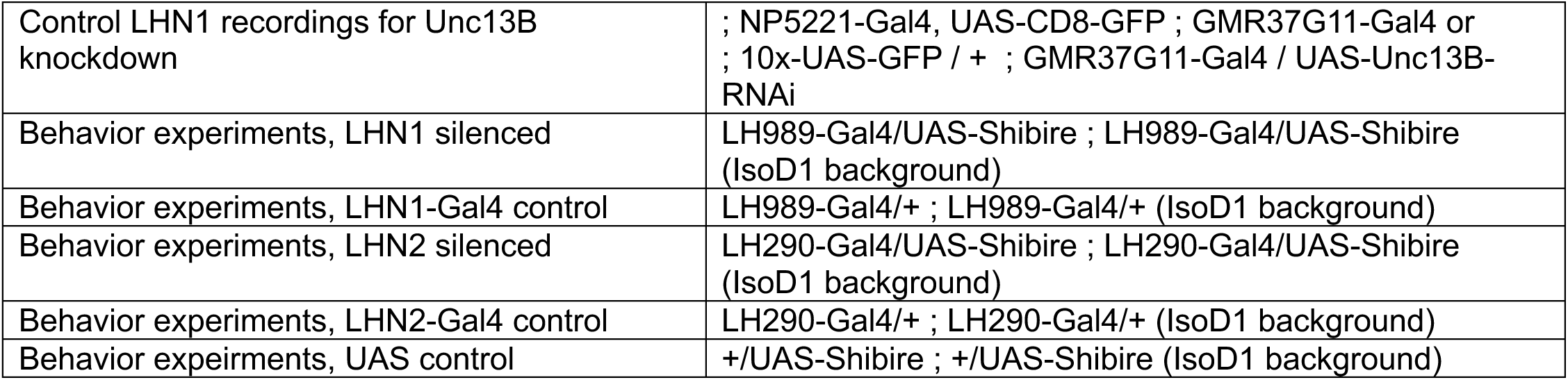

### METHOD DETAILS

#### Fly preparation for imaging and electrophysiology

For odor coding experiments, each fly was cold-anesthetized, positioned into a small holder made of stainless steel shim stock (0.001” thick), and affixed into position using paraffin wax such that the antennae were under the foil and dry while the head and brain was bathed in external saline. The external saline contained (in mM): 103 NaCl, 3 KCl, 5 N-tris(hydroxymethyl) methyl-2-aminoethane-sulfonic acid, 8 trehalose, 10 glucose, 26 NaHCO_3_, 1 NaH_2_PO_4_, 1.5 CaCl_2_ and 4 MgCl_2_ (osmolarity adjusted to 270–275 mOsm). A small window in the top of the head capsule was dissected using electrolytically sharpened tungsten wires and fine forceps. Fat, air sacs, and trachea were removed from above the brain. Fine forceps were then used to gently remove the perineurial sheath only above the area of the brain housing the target somata. Large-bore cleaning pipettes were then used to remove residual glia and interfering somata to gain clear access to target somata. External saline was bubbled with 95% O_2_ and 5% CO_2_ and reached an equilibrium pH of 7.3. Saline was superfused continuously over the brain during recording.

#### Voltage Imaging

*In vivo* voltage imaging of LHNs (Figure 1B and S2G) was performed on an upright epifluorescence microscope (Scientifica) and placed under a 40x 0.8NA water-immersion objective lens (Olympus). Arclight^79^ was excited with a 470 nm LED at 2.4% power (CoolLED pE-100), yielding 0.282 mW at the sample. Arclight fluorescence emission was collected with a scientific CMOS camera (Hamamatsu C1140-42U30), operating under HCImageLive software at an acquisition rate of 100 frames per second.

#### Calcium Imaging

In vivo calcium imaging of PNs (Figure S1) was performed on 2-photon laser scanning microscope (Scientifica MP-2000) using a 20x 1.0NA water-immersion objective lens (Olympus). A Titanium-Sapphire laser was tuned to 920nm to excite GCaMP6f, and fluorescence emission was collected with a GaASP PMT, controlled by ScanImage software (MBF Bioscience). Images were collected at an acquisition rate of 16.9 frames per second. Measurements from multiple z-slices through the antennal lobe were collected sequentially for each fly. DM1 and DM4 glomeruli were identified based on established landmarks^80^. No substantial activity was observed outside of DM1 and DM4 in these experiements.

#### Electrophysiology

Recordings were obtained using an Olympus BX51 upright microscope with 40X water immersion objective. One PN or LHN was recorded per fly, except for paired recordings. Patch-clamp electrodes were filled with an internal solution of (in mM): KCH_3_SO_3_H 160, HEPES 10, MgATP 4, Na_3_GTP 0.5, EGTA 1, biocytin hydrazide 13 (pH = 7.3, adjusted to 265mOsm). For paired recordings, EGTA was omitted from the internal solution in the PN pipette, except when explicitly stated otherwise. All other recordings included 1mM EGTA. Patch pipettes were made from borosilicate glass (Sutter; 1.5-mm outer diameter, 0.86-mm inner diameter) and were pulled and pressure polished^81^ to create a relatively long taper with final pipette tip opening of about 0.75μm in diameter. This enabled high seal resistances (>50GΩ), which helped keep leak currents negligible in LHNs, which typically have input resistances of 2-5 GΩ. Spike amplitudes in LHN recordings were typically ∼5-10mV and spike amplitudes in PN recordings were typically ∼15-25mV. Recordings were obtained with Axopatch 200B model amplifiers in current clamp mode with CV-203BU head stages, digitized at 10-30Hz with an analog-to-digital converter (National Instruments), and saved to disk using the MATLAB data acquisition toolbox. Recordings were not corrected for a liquid junction potential of ∼13mV^57^.

Paired PN-LHN recordings were conducted with the fly head rotated 90 degrees so that one eye was directed up. The eye and optic lobe were removed, along with the ipsilateral antenna to improve physical access to PNs. This configuration was necessary for recording pipettes to access both PN and LHN2 cell bodies because they are nearly antipodal to each other in the anterior-posterior and dorsal-ventral axes but are both lateral. Although DM1 and LHN1 cell bodies are in locations that are compatible with preserving both antennae^13^, we used the same head orientation for both sets of paired recordings here to minimize differences between them.

Current was applied to DM1 via the patch clamp amplifier (in current clamp mode) in 100 msec pulses at 5Hz (i.e., 100 msec intervals between pulses). A single exception was one DM1-LHN2 recording (with EGTA) where pulses were 60msec. Pulse amplitude was 100-300pA, determined empirically for each PN to evoke ∼6-8 spikes per pulse. The current amplitude was kept the same after dialysis, except for one pair where the amplitude was slightly increased. For square wave current injections into the PN, the series resistance was estimated from the fastest time constant of depolarization and subtracted from recorded PN voltage waveforms. Series resistance correction was imperfect during brief transient windows (<10msec in duration) at current onset and offset, so voltage data were interpolated from surrounding values during these times.

For all paired experiments, the recording from the LHN was established first, and then the DM1 recording was established. This allowed immediate measurements of synaptic potentials in the paired configuration before PN dialysis. Because the contralateral antenna remained intact, odor stimulation could still drive responses. In one DM1-LHN2 paired recording (with EGTA), we used a solid 2 seconds of ethyl acetate (1e-6× concentration) instead of pulsed current injection. This pair is included in Figure 4G-J (using an identical 200msec analysis window), but not in Figure 4E,F, because the temporal stimulus profile was different.

We note that 1mM EGTA is commonly included in internal saline recipes in *Drosophila* experiments. While EGTA does not appear to affect a cell’s integration of inputs and spike generation (as measurable from the soma), it clearly disrupts some synaptic outputs. Thus, we recommend omitting EGTA from internal saline during any patch-clamp recording where perturbation to its synaptic outputs is not desired (e.g., during simultaneous measurement of components of downstream network activity or of behavior).

Following most LHN recordings, brains were processed for immunohistochemistry exactly as described previously^14^, and imaged with a Zeiss LSM 880 confocal microscope to visualize the biocytin fill. All filled cells of each LHN type were morphologically alike to a degree expected of a single cell type^12,82^. No recordings were omitted based on this recovered morphology.

#### Tethered walking behavior

Individual flies were anesthetized on ice in a custom-built jig to hold the fly in an appropriate position for tethering. A small amount of UV glue (BONDIC) was applied on the tip of a ½” long 30-gauge needle (BD) which was then carefully lowered onto the dorsal thorax of the fly at an angle of 45-60°. The glue was cured with UV light for 40 seconds to firmly attach the dorsal thorax to the needle tip. The base of the needle was then mounted to a platform above the air-supported ball.

The hollow plastic ball had a radius of 3.175 mm. To increase contrast for automated rotation tracking, the ball was painted with white acrylic paint to provide a homogenous background, on which black patches were subsequently drawn with black acrylic paint to provide a random spatial pattern. The ball was floated from a semi-spherical concave metal socket by a constant air stream (550 mL/min, which was sufficient for the ball to spontaneously roll while floating). The metal socket and the airflow system were mounted on a micromanipulator to precisely position the ball under the tethered fly. A 3D-printed square plastic frame was added at the base of the metal socket to provide a readily discernible reference frame for automated tracking. An IR-sensitive digital camera (Basler ace acA1300-200um) with long working distance lens (Infinity Optics) was installed facing the fly’s left side such that the square plastic frame, the walking fly, and exposed subset of the ball were all within its field of view. Videos were recorded at 30 frames per second.

The entire setup was mounted on a vibration isolation table and enclosed in a large cage with black cardboard walls and ceiling to block any visible light and to achieve adequate insulation for thermogenetic manipulation (see below). Inside the enclosure, an infrared LED (810 nm Thorlabs M810L4) was installed above the setup to illuminate the ball for the camera without being visible to the fly. Odors were delivered to the front of the fly in an open loop configuration (i.e., the fly’s walking velocity did not affect odor delivery).

Using the micromanipulator, the ball position was carefully adjusted so that the fly’s legs were in contact with the ball and its ventral thorax roughly horizontal. The fly was then given sufficient time (typically 15-30 minutes) to recover from anesthesia and acclimate to the ball (i.e., when it began to initiate spontaneous naturalistic walking bouts, which were used as a filtering criterion for proceeding with the experiment). For the flies that failed to acclimate (i.e. did not readily walk) and/or showed signs of agitation (such as frantic leg flailing and abdomen bending), the experiment was aborted and excluded from all analysis.

To silence neurons using *Shibire^ts^*, the entire enclosure containing the fly, cameras, and odor delivery device, was heated using a ceramic infrared heat lamp equipped with a remote temperature sensor and feedback control system (Wuhostam 50W). This maintained a steady temperature inside the enclosure of ∼30.5°C. The lamp was installed facing the fly at an angle of 45 degree from above. The sensor, placed next to the tethered fly, continuously read the ambient temperature, and the feedback control system automatically corrected for any deviation of the ambient temperature from the preset target temperature by controlling the lamp output. Behavioral experiments for all genotypes (including controls) were conducted at the same 30.5°C temperature. As a positive control to validate that the *Shibire^ts^* construct functioned as intended, UAS-Shibire^ts^ (on IsoD1 WT background) was crossed with the pan-neuronal driver line nSyb-Gal4. Indeed, at 30.5°C, these flies were completely paralyzed.

#### Odor delivery and stimulus design

A clean air stream (1360 mL/min for electrophysiology and imaging, 360 ml/min for behavior) was filtered through activated carbon and directed to the fly through a carrier tube (**Figure S1**). Separate air streams of 12 ml/min were directed under the control of a solenoid valve (The Lee Company, model LHDA1231415H) into the headspace of 2mL vials (Thermo Scientific, National C4011-5W) containing odors. The odor streams joined the carrier stream 9-11 cm from the end of the tube. The experiments reported in Figures 1, 2, 3, 5, and 6F-J used a single valve per odor vial and predominantly used Tygon tubing. The experiments reported in Figure 6A,D,E and 7 used two valves per odor vial and predominantly used PTFE tubing. Schematics of these odor delivery devices are provided in Figure S1A. In some experiments, a digital camera (Basler ace acA1300-200um) was set up facing the lateral side of the prep, to facilitate uniform positioning of the carrier tube relative to the fly antennae across experiments.

For electrophysiology and imaging, low concentrations of ethyl acetate and methyl acetate (both at 1e-6×) were prepared by serial dilution in either paraffin oil or water. For behavior, apple cider vinegar was diluted in water to either 10% of 50% concentration. To ensure sufficient headspace in each odor vial, the final volume of odor dilutions was 0.5 mL.

The timing of valve opening and closing (and therefore odor delivery) was controlled by a custom MATLAB script. For experiments reported in Figures 1, 2, 3, 5, and 6F-J, individual odor pulses were 40 msec in duration. For experiments reported in Figures 6A,D,E and 7, individual odor pulses were 150 msec in duration. The longer duration was necessary because the two-valve odor delivery system required more time for odor to reach the fly. Because two odors were simultaneously connected to the carrier stream and were independently controllable, we alternated odor identity from trial to trial and interleaved odor pulse frequencies, to remove the effects of any slow changes in neural responses or behavior over the course of each experiment.

For characterization of unadapted odor tuning curves for each neuron type, odors were delivered in 40msec pulses at 0Hz (i.e., spontaneous activity) 1Hz, 2.5Hz, 5Hz, and 10Hz, each following at least 10 seconds without any odor stimulation. 10 pulses were delivered for 1Hz stimulation, 20 pulses for 2.5Hz, and 30 pulses for 5 and 10Hz stimulation. To measure adapted odor tuning curves, 20 seconds of pulsed odor stimuli were delivered at “prior” frequencies of 1, 2.5, or 5Hz, and then odor pulse frequency was immediately changed to the “current” frequency of 1, 2, 2.5, 3, 3.5, 5, 7.5, or 10Hz for 10 seconds. Current frequencies of 2, 3, 3.5, and 7.5 were not tested for every prior frequency. For cross-adaptation experiments (**Figure S4**), the adapting frequency was always 5Hz (of ethyl acetate).

Valve switching introduced brief and small electrical artifacts in electrophysiology voltage traces. These were most evident for the high input resistance neurons (LHNs) where small noise currents cause proportionally larger voltage deflections. For display purposes only, we removed these artifacts by blanking the voltage for 10-12msec after each valve switching event, and linearly interpolating to fill the gap.

### QUANTIFICATION AND STATISTICAL ANALYSIS

#### Connectome analysis

Synaptic connectivity data were obtained from the Hemibrain connectome (version 1.2.1) using custom Python scripts and the neuprint-python API. Synapse counts between all canonical cholinergic uniglomerular PNs (those in the adPN or lPN lineages) and each instance of LHN1 and LHN2 were retrieved with “fetch_adjacencies.” For analysis of PN outputs (**Figure S1A**), raw synapse counts were normalized by total PN output synapses. For analysis of LHN inputs (**Figure S1B**), raw synapse counts were normalized by the total number of input synapses on each LHN instance. Reciprocal connections onto LHN1 (**Figure S7B**) were identified similarly, and raw synapse counts were normalized by total synapse count of the recipient neuron. Synapse location and cellular morphology data (**Figure S5**) were obtained from the same connectome using custom R scripts using the “hemibrain_extract_synapses” and “hemibrain_read_neurons” functions of the natverse toolbox^83^. Euclidean and geodesic distances between all pairs of synapses on PN axon arbors were computed using the TREES toolbox for MATLAB^84^.

#### Spike identification, analysis of spike rates, and analysis of membrane voltages

Prior to spike detection, raw voltage traces were highpass filtered with a second-order digital Butterworth filter with cutoff frequency of 5Hz and then lowpass filtered with a second-order digital Butterworth filter with cutoff frequency of 500Hz. Spikes were detected as negative threshold crossings of the second derivative of the filtered voltage trace. The threshold was adjusted for each recording. Because recording conditions can change slightly over time, every threshold crossing was inspected to remove artifactual false positive crossings. Artifactual false negatives were manually added back in. Most of the false positive crossings were the result of valve switching artifacts described above (to ensure every spike was detected, spike detection was performed without removal or interpolation of these artifacts).

Peristimulus time histograms (PSTHs) were computed by binning spike times into 50msec bins. All repeated trials for each stimulus were combined into a single average PSTH for that neuron and stimulus. For all unadapted odor-tuning curves and cellular input-output functions, the transient responses for each stimulus were computed as the average spike rate in the first 1 second of the response. For adapted tuning curves, the transient responses were computed as the average spike rate in the first 1 second after switching to the new odor pulse frequency. Because our odor delivery system required about 100msec for odor to flow from the vial to the fly (**Figure S1**), we shifted each response window (relative to valve timing) by 100msec to account for this.

Trough voltages (**Figure 6D**) were measured as the average membrane voltage in 50msec windows starting 750msec after the response to each odor pulse (i.e., the 50msec just prior to the depolarizing response to the next odor pulse). Normalized odor responses in **Figure 6E** were computed over 150msec windows starting 180msec after valve opening. The longer latency between valve opening and the start of the odor response in these experiments was due to the use of a slightly different odor delivery device (**Figure S1**).

#### Curve fitting to odor tuning curves and cellular input-output functions

Unadapted odor tuning curves (functions relating odor pulse rate to spike rate) and cellular input-output functions (relating PN spike rates to LHN spike rates) were fit with the saturating hyperbolic ratio function^24,85^

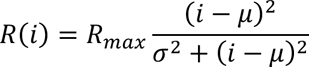

where the variable *i* is the input (PN spike rate or odor pulse rate) and *R* is the output (PN or LHN spike rate). *R_max_* represents the saturation value of the function, *μ* represents the subtractive input offset, and *σ* determines the divisive input gain; all three parameters were fitted to data. Divisive adaptation was modeled by adjusting *σ* (a standard model of divisive gain control^86^) and subtractive adaptation was modeled by adjusting *μ*. Fits to adapted data were carried out by freezing R_max_ at the value fitted to unadapted data and then refitting both both *μ* and *σ*. For all adapted fits, *R* was set to zero for *i* < *μ*, and *μ* and *σ* were each constrained to be equal or larger than the corresponding fit to unadapted data (to avoid spurious multiplicative or additive modulation fits).

Measures of baseline adaptation and nonlinearity sharpness were computed by linear regression to spike rates for each LHN individually (as a function of average DM1 spike rates). Baseline adaptation was computed for spike rates corresponding to odor stimulus frequencies between 0 and 5Hz. The nonlinearity sharpness was the difference in gain between linear regressions for odor stimulus frequencies between 0 and 5Hz, and between 5Hz and 10Hz. The gain ratio (Figure 6I) was computed as the ratio of the gain for linear regression for stimulus frequencies between 0 and 10Hz for adapted responses and the gain for linear regression for stimulus frequencies between 0 and 5Hz unadapted responses. The stimulus domains used for regression was different in order to only capture the linear regime of responses (in the unadapted case, the 10Hz stimulus began to engage the saturating nonlinearity).

#### Synapse analysis

Unitary excitatory postsynaptic potentials (uEPSPs) were computed from the membrane voltage in a brief time window surrounding each DM1 PN spike time (starting 1msec prior to the PN spike until 10msec after the PN spike). If the LHN generated a spike during this time window, the uEPSP waveform was omitted from all further analysis. In a few cases, LHNs were gently hyperpolarized by 5-10mV with tonic current injection to reduce likelihood of spiking.

For the analysis of EGTA effects, pre-dialysis measurements were all taken within 8 minutes of establishing the PN whole cell patch clamp configuration (mean: 3.3 minutes). Post-dialysis measurement were all taken after at least 12 minutes of establishing the PN whole cell patch clamp configuration (mean: 33.5 minutes). Synaptic responses remained stable following dialysis.

We used quantal theory to estimate the initial release probability and quantal content of PN-LHN synapses (**Figure 3E**). Quantal theory relates the mean synaptic current amplitude to the number of release sites (*N*), probability of release at each site (*P*), and the quantal size of each release event (*Q*)^87^

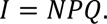

Assuming a binomial model^88^, the variance of the synaptic current amplitude (over repeated trials) is

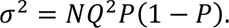

In *Drosophila*, the number of release sites estimated by quantal theory^58^ closely matches the number of physical synapses measured with electron microscopy^89^. Thus, we used synapse counts from the Hemibrain connectome to estimate the mean *N* for each synapse type (43.0 ± 8.4 for DM1-LHN1, and 18.8 ± 4.4 for DM1-LHN2; incidentally, our biocytin fills indicated that this LHN1 sample only included the PD2a1 subtype, so estimates were taken only from that subtype in the Hemibrain)^75^. This allows *P* and *Q* to be calculated from the equations above using experimental measurements of the mean and variance of EPSP amplitude. Because quantal theory predicts the amplitude of synaptic current instead of voltage (our experiments only measured voltage), we estimated EPSC amplitude by diving EPSP amplitude by the average input resistance for each cell type (2.7 ± 0.5 GΩ for LHN1, 4.1 ± 0.9 GΩ for LHN2). We only analyzed the first EPSP in the first pulse of each trial and verified that none of these responses evoked spikes in the LHN.

#### Contrast and prediction error coding

Comparison of LHN coding of proportional changes (contrast) and absolute differences (temporal prediction error) in odor frequency were performed by either dividing the test frequency by the adapting frequency (contrast) of subtracting the adapting frequency from the test frequency (prediction error) prior to plotting the odor tuning curves in **Figures 1I,J, 5I**, and **6J**). Data for each of the three adapting frequencies (1Hz, 2.5Hz, 5Hz) are shown separately (except in **Figure 5I**, where they are pooled), but unadapted data are not shown, since the proportional change in frequency would be undefined in this case. All data for 2.5Hz and 5Hz adaptation are shown (and included in the linear fits). The two highest test frequencies for 1Hz adaptation are omitted because these started to engage the saturating nonlinearity, which we did not consider in this analysis. Linear models were fit to data for each adapting frequency separately, but only nonzero responses were included for fits for LHN2, again to focus on the linear spike rate regime. The ANCOVA test was used to determine whether a single model relating LHN spike rate to proportional changes or absolute differences in odor pulse rate could be rejected.

Knockdown of Unc13B was performed by expressing Unc13B-RNAi in DM1 PNs. Because of genetic constraints, this manipulation also knocked down Unc13B in LHNs (since both PNs and LHNs were targeted with the Gal4/UAS system). Since Unc13 occurs only at presynapses^40^, our manipulation should not have any direct LHN effects in our experimental configuration. In agreement with this specificity, we did not see any effects of Unc13B knockdown on LHN responses to methyl acetate (which does not engage DM1 PNs) or on intrinsic LHN2 excitability (**Figure S6**). In addition, in some experiments we expressed Unc13B-RNAi only in LHN2 (and not in DM1), which did not affect on ethyl acetate responses. These data are included in the control datasets in **Figure 5C-E** and in **Figure S6**.

#### Behavior analysis

The tethered fly’s fictive locomotion velocity in response to each odor stimulus was recorded in 45 second videos using the digital camera. The displacement of the black and white pattern on the ball surface was tracked frame by frame using FicTrac^90^ to build a global map of the ball surface and compute the instantaneous angular velocity of the ball along each rotational axis. These rotational measures were then used to deduce the fly’s total instantaneous walking speed (independent of direction, since the odor tube was fixed in position in front of the fly). Three (sometimes four) trials were conducted for each odor stimulus in each fly. Extracted data were averaged across trials for each condition, then across all flies belonging to the same genotype. Because the baseline walking speed was variable from fly to fly, we normalized the speed of each fly to its baseline, to compute the fold-change in walking speed.

---

**Figure S1 (related to Figure 1):**
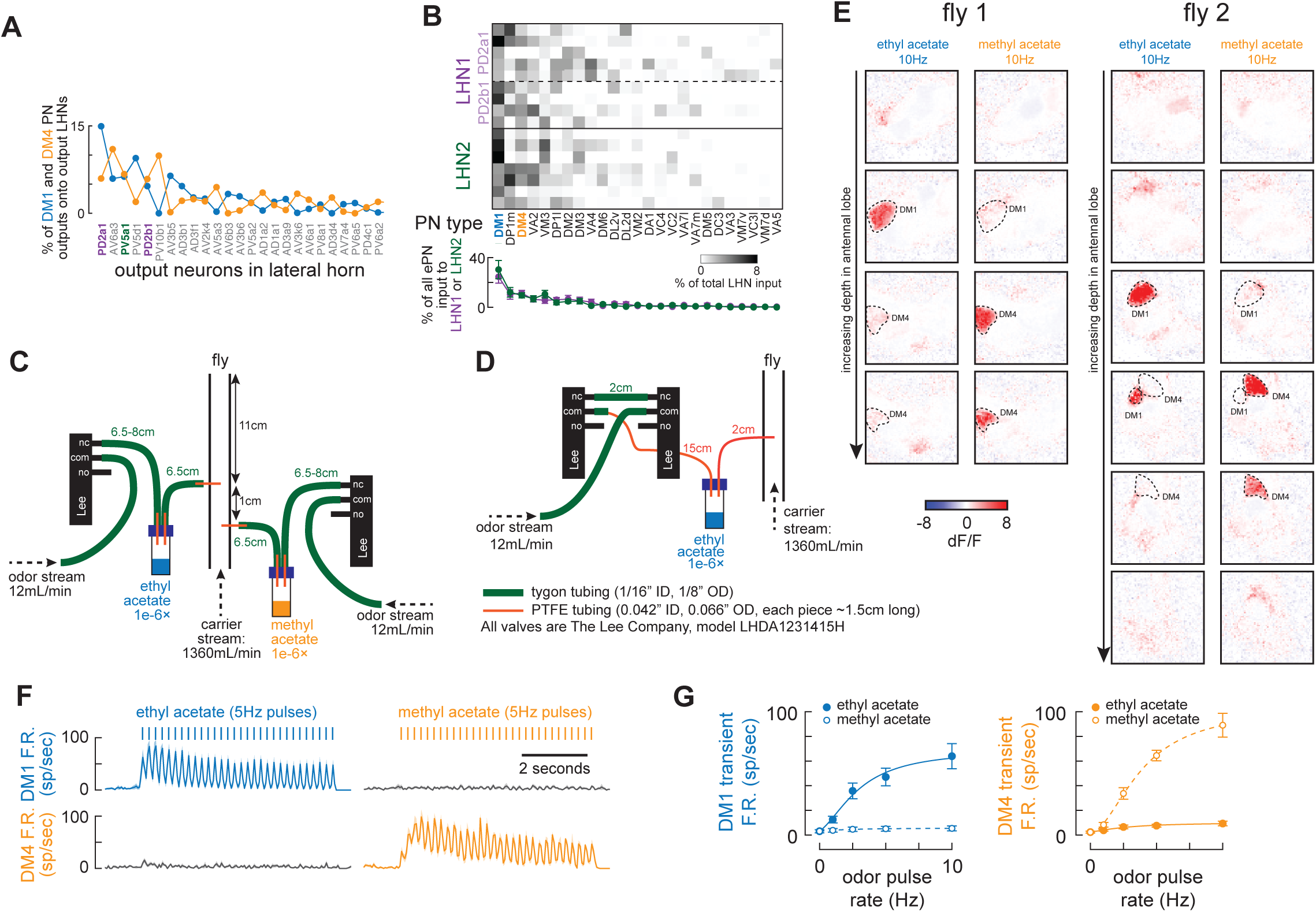
Connectivity, odor delivery, and private odor validation. **(A)** Average distribution of synapses onto output LHN types from the DM1 and DM4 PNs (as a fraction of all synapses from each PN that target output LHNs). **(B)** Top: input to each instance of LHN1 and LHN2, for the top 25 cholinergic uniglomerular PN inputs (as a fraction of all input to each LHN). Bottom: average distribution of synapses onto LHN1 and LHN2 types from the top 25 cholinergic uniglomerular PNs (as a fraction of all cholinergic uniglomerular inputs per LHN type). **(C)** Schematic of odor delivery device used for all experiments except for those in Figure 6A,D,E and in Figure 7. **(D)** Schematic of odor delivery device used for experiments in Figure 6A,D,E and Figure 7. This design used an extra valve so that the headspace in the odor vial was always vented away from the carrier stream. This was designed for the use of higher concentration odors to prevent odor leaking into the carrier stream. However, none of the odors used in this study were high enough concentration to cause leakage. The only functional difference between the designs in (C) and (D) is that (D) has a longer latency for odor to reach the fly. Thus, for the data reported in Figure 6A,D,E and in Figure 7, valves were opened for 150msec instead of 40msec. nc = normally closed. no = normally open. com = common. Note that a lower carrier stream was used for behavioral experiments in Figure 7. **(E)** 2-photon calcium imaging of GH146-Gal4 / UAS-GCaMP6f in several planes in the antennal lobe in 2 different flies, in response to 10Hz ethyl acetate or 10Hz methyl acetate (the highest intensity odors used for physiology). Ethyl acetate is specific for DM1 and methyl acetate is specific for DM4, as seen with electrophysiology. No substantial calcium response in any other glomerulus is observed, indicating that our stimuli are private, consistent with previous studies that have used these odors. **(F)** Mean (± s.e.m.) PSTHs of DM1 (top, n = 8) and DM4 (bottom, n = 5) spike rates in response to 5Hz ethyl acetate (left) and methyl acetate (right). Spike rates are determined from whole cell patch clamp electrophysiology. **(G)** Tuning curves of initial transient spike rates (mean ± s.e.m., computed within the first 1 second of response) for ethyl and methyl acetate in DM1 (n = 8) and in DM4 (n = 5). The lack of substantial responses for the off-target odor in each PN type further validates the privacy of these odor stimuli.

**Figure S2 (related to Figure 1):**
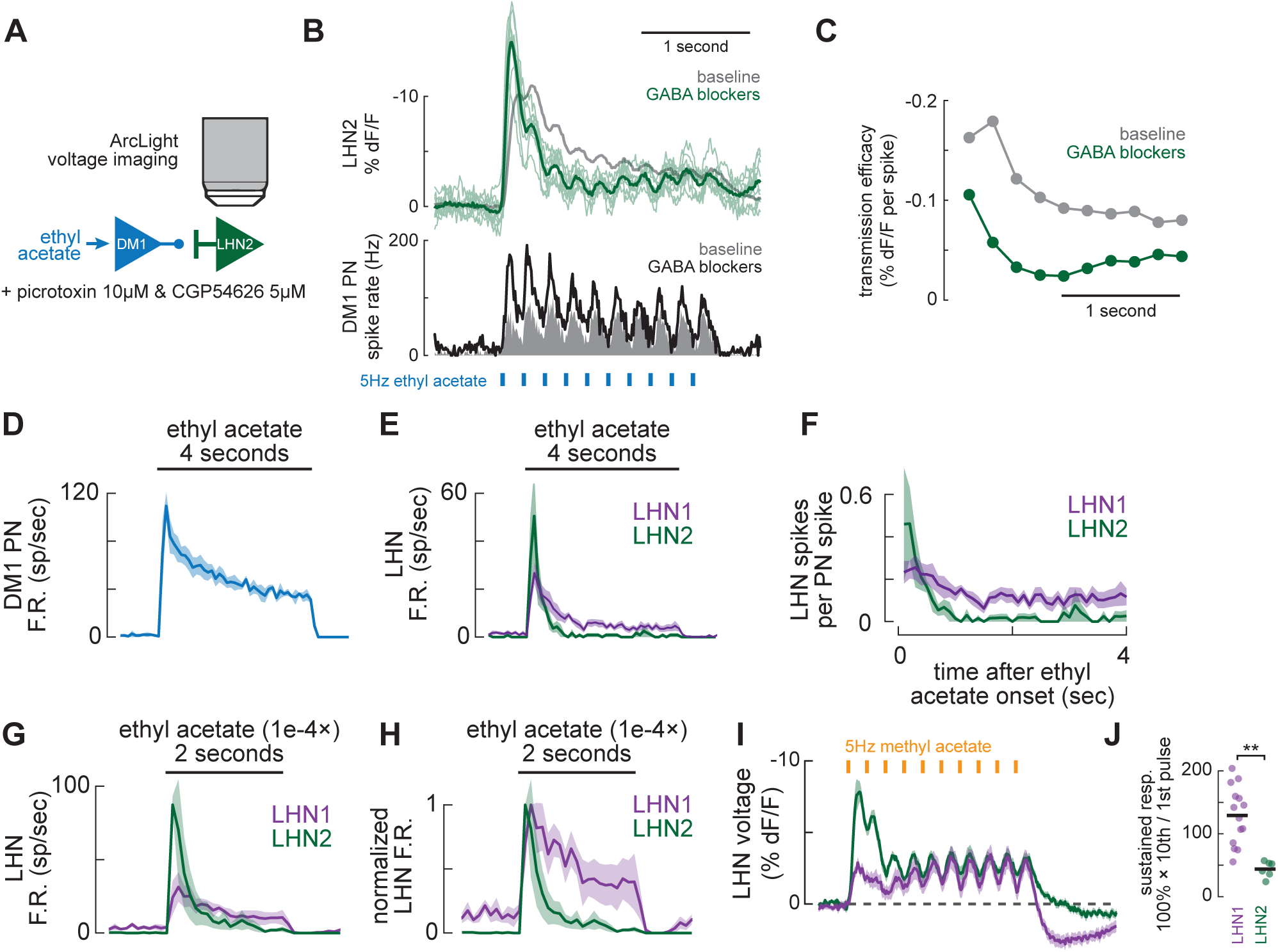
LHN2 adaptation does not require synaptic inhibition and differences in LHN dynamics are not stimulus specific. **(A)** Schematic of experimental test of the role of synaptic inhibition in LHN2 dynamics. In each experiment, baseline odor responses were recorded, picrotoxin and CGP54626 were bath-applied to block GABA-A and GABA-B receptors, and then odor responses were recorded again. Because bath application also affects PN responses to odors, we also measured PN spike rates during identical pharmacological manipulation. **(B)** Top: comparison of responses to 5Hz ethyl acetate in LHN2 before and after pharmacological blockade of GABA receptors. With blockers, adaptation to odor pulses becomes slightly stronger (thin green lines are single recordings, thick green line is the mean) than the mean response in control saline (gray). Bottom: GABA blockers also increase DM1 PN adaptation, mostly by increasing responses to the first few odor pulses. **(C)** Voltage responses per odor pulse in LHN2 normalized by the number of DM1 spikes per pulse. Each point is the mean response to each odor pulse. Blocking GABAergic inhibition does not dramatically change the dynamics of the “transmission efficacy” relating PN spikes to LHN voltage. **(D)** Mean (± s.e.m.) PSTHs of DM1 (n = 3) in response a 4-second solid pulse of ethyl acetate (at 1×10-6 concentration), showing that responses are sustained, but adapt modestly, similar to pulsed odor stimulation. **(E)** Mean (± s.e.m.) PSTHs of LHN1 (n = 8) and LHN2 (n = 3) in response to the same 4-second solid pulse of ethyl acetate as in panel A, showing sustained activity in LHN1, but not in LHN2, similar to pulsed odor stimulation **(F)** Mean (± s.e.m.) LHN spikes per PN spike during the same 4-second solid pulse of ethyl acetate (same data as in panels A and B). Responses are binned into 100msec bins. Both LHN types adapt, but LHN2 adapts more strongly than LHN1, similar to our main observations with pulsed stimuli. **(G)** Mean (± s.e.m.) PSTHs of LHN1 (n = 4) and LHN2 (n = 3) responses to a 2-second solid pulse of ethyl acetate at 1×10-4 concentration, which activates many glomeruli, including DM1 and DM4. **(H)** Same as (D), but with PSTHs normalized by their peak value. **(I)** Mean (± s.e.m.) voltage change (ArcLight fluorescence) of LHN1 (n = 10) and LHN2 (n = 12) in response to 5Hz pulses of methyl acetate (at 1×10-6 concentration, which privately activates the DM4 glomerulus; Figure S1). Voltage scale is inverted because fluorescence decreases with voltage increases.

**Figure S3 (related to Figures 1 and 2):**
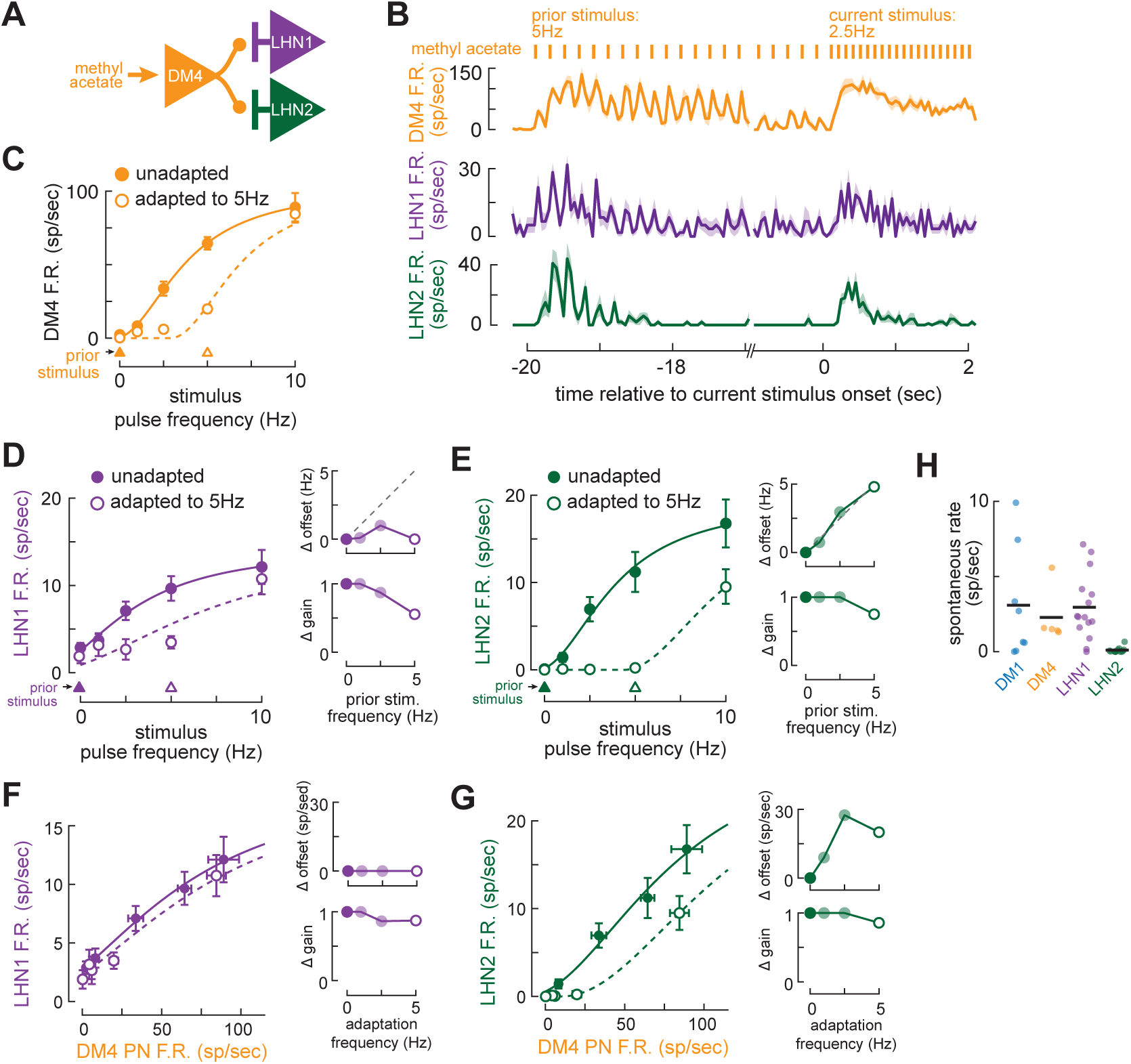
Divergent processing of DM4 inputs to LHN1 and LHN2. **(A)** Schematic of divergence from the DM4 PN onto LHN1 and LHN2. **(B)** Mean (± s.e.m.) PSTHs of DM4 (top, n = 5), LHN1 (middle, n = 6), and LHN2 (bottom, n = 5) in response 20 seconds of 5Hz pulses of methyl acetate, when is then switched to 10Hz. **(C)** Tuning curve of mean (± s.e.m.) DM4 PN (n = 4-5) spike rates to a range of methyl acetate pulse frequencies without adaptation (solid circles) and adapted to a prior frequency of 5Hz (open circles). Curves are fit with the same procedure and model as for DM1 PNs (STAR Methods). As with DM1, adapted responses were best fit with a pure subtractive shift, but responses to low stimulus frequencies were notably underpredicted by this model. **(D)** Left: tuning curves for mean (± s.e.m.) LHN1 (n= 6-16) spike rates to a range of methyl acetate pulse frequencies without adaptation (solid circles) and after adaptation to a prior frequency of 5Hz methyl acetate (open circles). Curves are fit with the same procedure and model as for ethyl acetate stimulation (STAR Methods). Right: offset and gain adjustments for the best fits to each adaptation frequency. **(E)** Same as (D) but for LHN2 (n = 5-12). **(F)** Left: input-output functions relating mean (± s.e.m.) DM4 spike rates to mean (± s.e.m.) LHN1 spike rates, without adaptation (solid circles), and after adaptation to a prior frequency of 5Hz methyl acetate (open circles). Data are the same as in panels C and D. Curves are fit with the same procedure and model as for ethyl acetate stimulation (STAR Methods). Right: offset and gain adjustments for the best fits to each adaptation frequency. **(G)** Same as (F) but for LHN2. Data are the same as in panels C and E. **(H)** Comparison of spontaneous spike rates of DM1, DM4, LHN1, and LHN2.

**Figure S4 (related to Figures 1 and 2):**
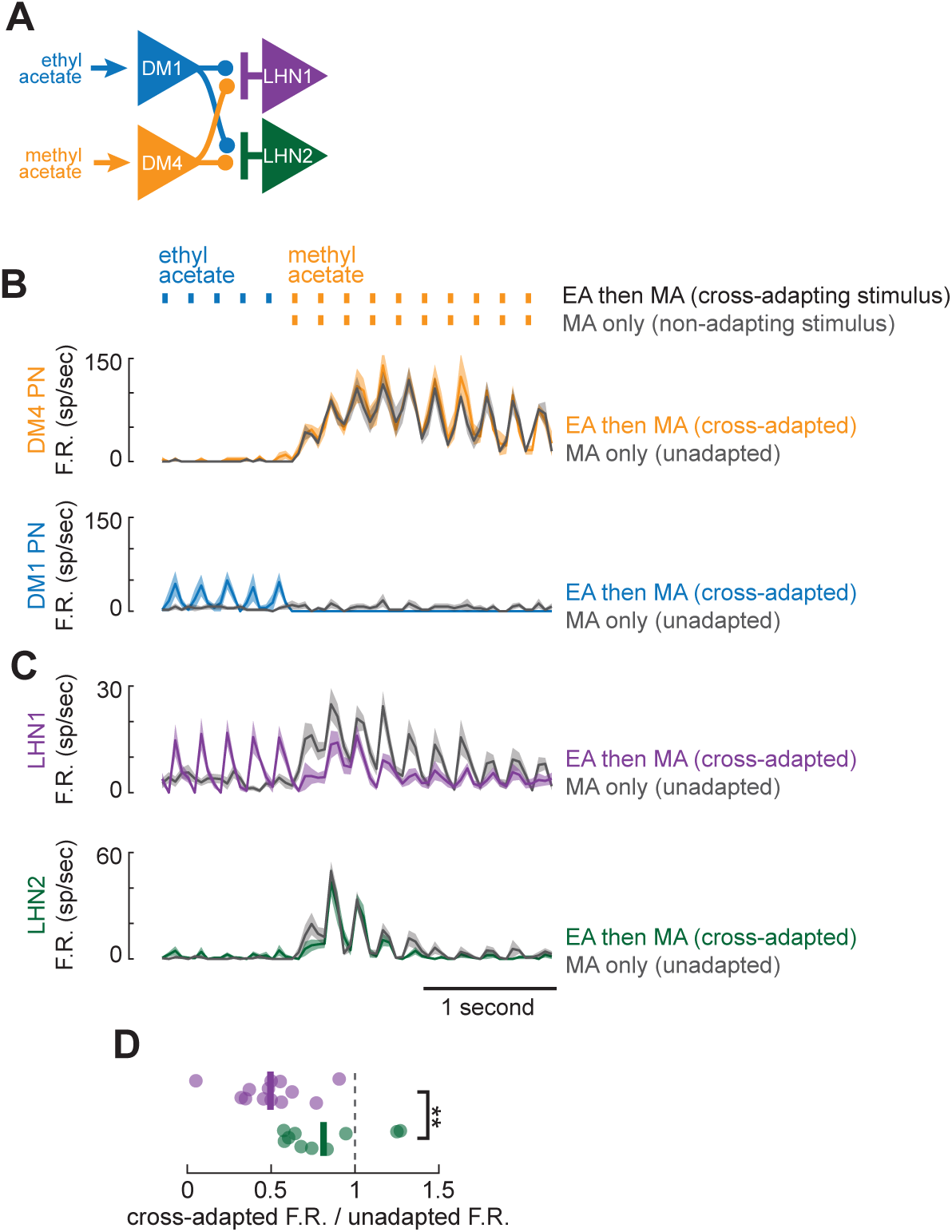
Adaptation in LHN2 is more input-specific than adaptation in LHN1. **(A)** Schematic showing the connectivity of DM1, DM4, LHN1, and LHN2 underlying the cross-adaptation experiment. Both DM1 and DM4 diverge onto LHN1 and LHN2. **(B)** Top: mean (± s.e.m.) responses of DM4 PNs (n = 3) to 5Hz ethyl acetate then 5Hz methyl acetate (orange) or to methyl acetate alone (gray). Bottom: mean (± s.e.m.) responses DM1 PNs (n = 4) to ethyl acetate then methyl acetate (blue) or to methyl acetate alone (gray). No evidence of cross adaptation is apparent in either PN type. These plots, and those in (C), only show the end of ethyl acetate stimulation, which had fully adapted by the time the odor switched to methyl acetate. **(C)** Top: mean (± s.e.m.) responses of LHN1 (n = 13) to 5Hz ethyl acetate then 5Hz methyl acetate (purple) or to methyl acetate alone (gray). Bottom: mean (± s.e.m.) responses of LHN2 (n = 10) to 5Hz ethyl acetate then 5Hz methyl acetate (green) or to methyl acetate alone (gray). **(D)** Normalized cross-adapted responses (cross-adapted F.R. / unadapted F.R.) are smaller in LHN1 than in LHN2 (** t-test p = 0.0043), indicating greater cross-adaptation in LHN1.

**Figure S5 (related to Figures 3 and 4):**
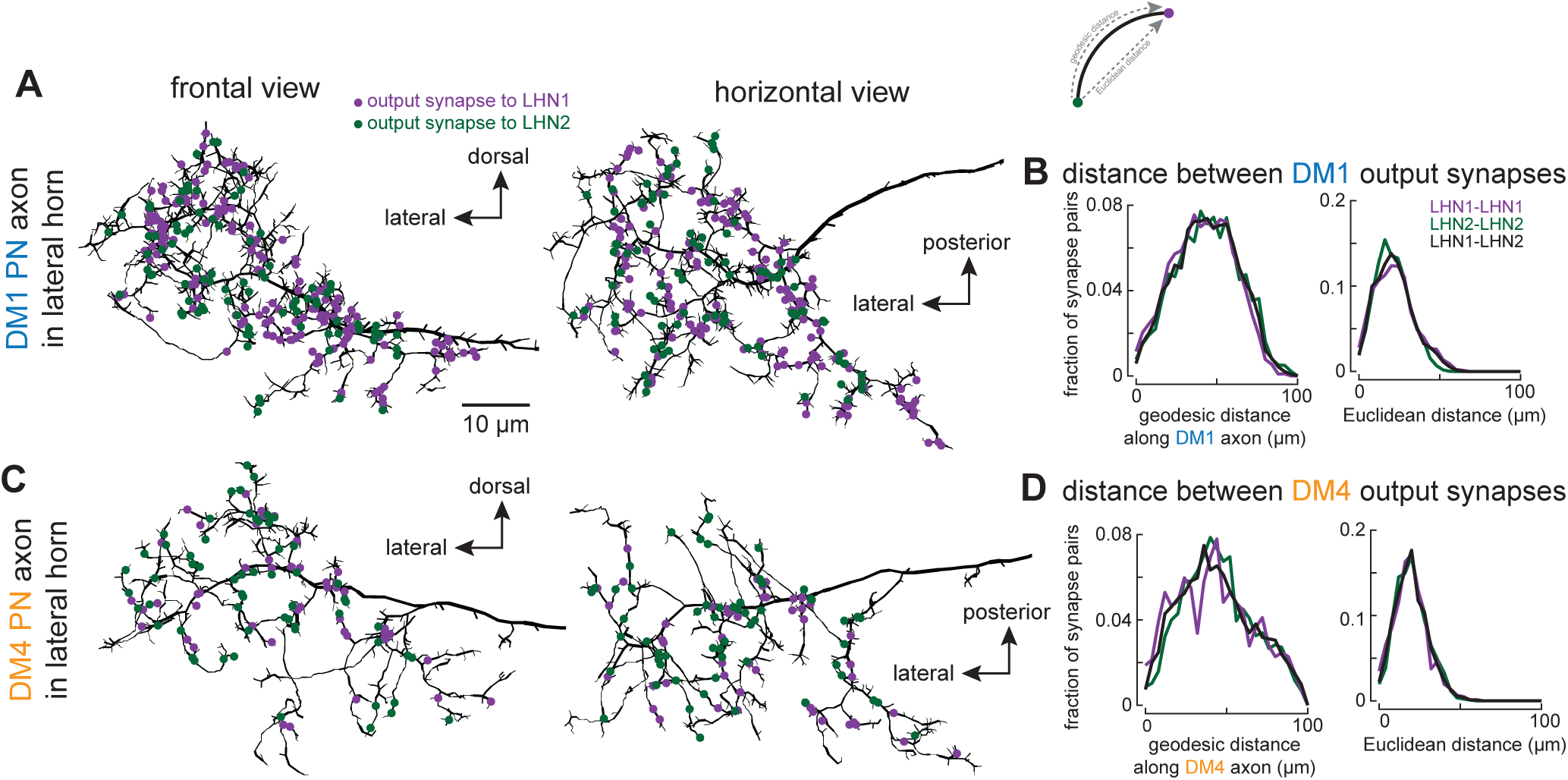
PN synapses onto LHN1 and LHN2 are not spatially organized. **(A)** Synapse locations on the DM1 PN axon in the lateral horn onto LHN1 (purple) and LHN2 (green). Frontal and horizontal are two different views of the same axon. There is no obvious spatial or branch-specific organization to synapse locations. **(B)** Distributions of geodesic (left) and Euclidean (right) distances between every pair of LHN1 and LHN2 synpases on the DM1 axon in the lateral horn. Geodesic distance is the distance along the neurite (see schematic above). There are no substantial differences between these distributions. **(C,D)** Same as (A,B) but for the DM4 axon.

**Figure S6 (related to Figure 5):**
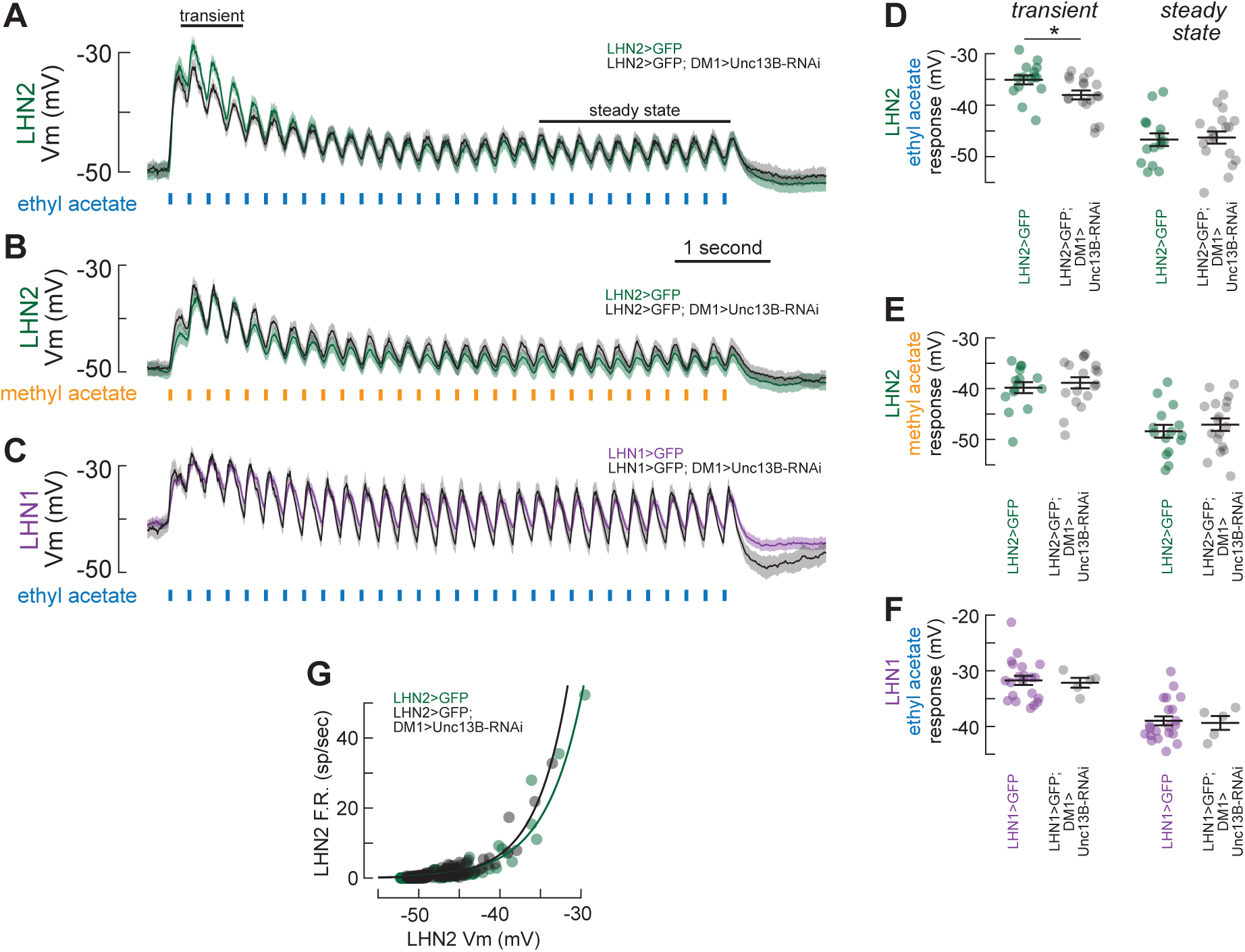
Unc13B knockdown is selective for DM1 inputs to LHN2. Genetic constraints required Unc13B knockdown in both DM1 and LHN2. Since Unc13B is only known to express in presynapses, it should not have any effects on LHN2 activity due to Unc13B-RNAi expression in LHN2 itself. If this is true, Unc13B-RNAi should only reduce the transient LHN2 depolarizations for ethyl acetate and not for methyl acetate. It should also not change the intrinsic excitability of LHN2. Finally, if its effects are selective for facilitating synapses, then it should have no effect on ethyl acetate responses in LHN1. **(A)** Mean (± s.e.m.) LHN2 membrane potential for control flies (green, n = 15) and flies with knockdown of Unc13B in DM1 (black, n = 18) in response to 5Hz ethyl acetate stimulation. **(B)** Mean (± s.e.m.) LHN2 membrane potential for control flies (green, n = 15) and flies with knockdown of Unc13B in DM1 (black, n = 18) in response to 5Hz methyl acetate stimulation. **(C)** Mean (± s.e.m.) LHN1 membrane potential for control flies (purple, n = 21) and flies with knockdown of Unc13B in DM1 (black, n = 5). **(D-F)** Mean transient and steady state responses (averaged of time ranges denoted by black bars in panel 1), for each of the three genetic comparisons in (A-C). * t test, p < 0.05. All other comparisons are not significant. **(G)** Comparison of the relationship between mean membrane voltage (as measured at the soma) and mean spike rate for LHN2 recordings in control conditions (green, n = 15) and with DM1>Unc13B-RNAi (black, n = 18). Each point is the mean response for each 250 msec bin for responses to ethyl acetate delivered at 10, 5, and 2.5Hz. Curves are single exponential fits. The requirement of Unc13B for elevated transient depolarizations in LHN2 (panel A,D) is thus likely due to specific effects at the DM1-LHN2 presynapse, and not any direct effects of Unc13B in LHN2. Moreover, these findings are consistent with Unc13B only operating at facilitating synapses.

**Figure S7 (related to Figure 6):**
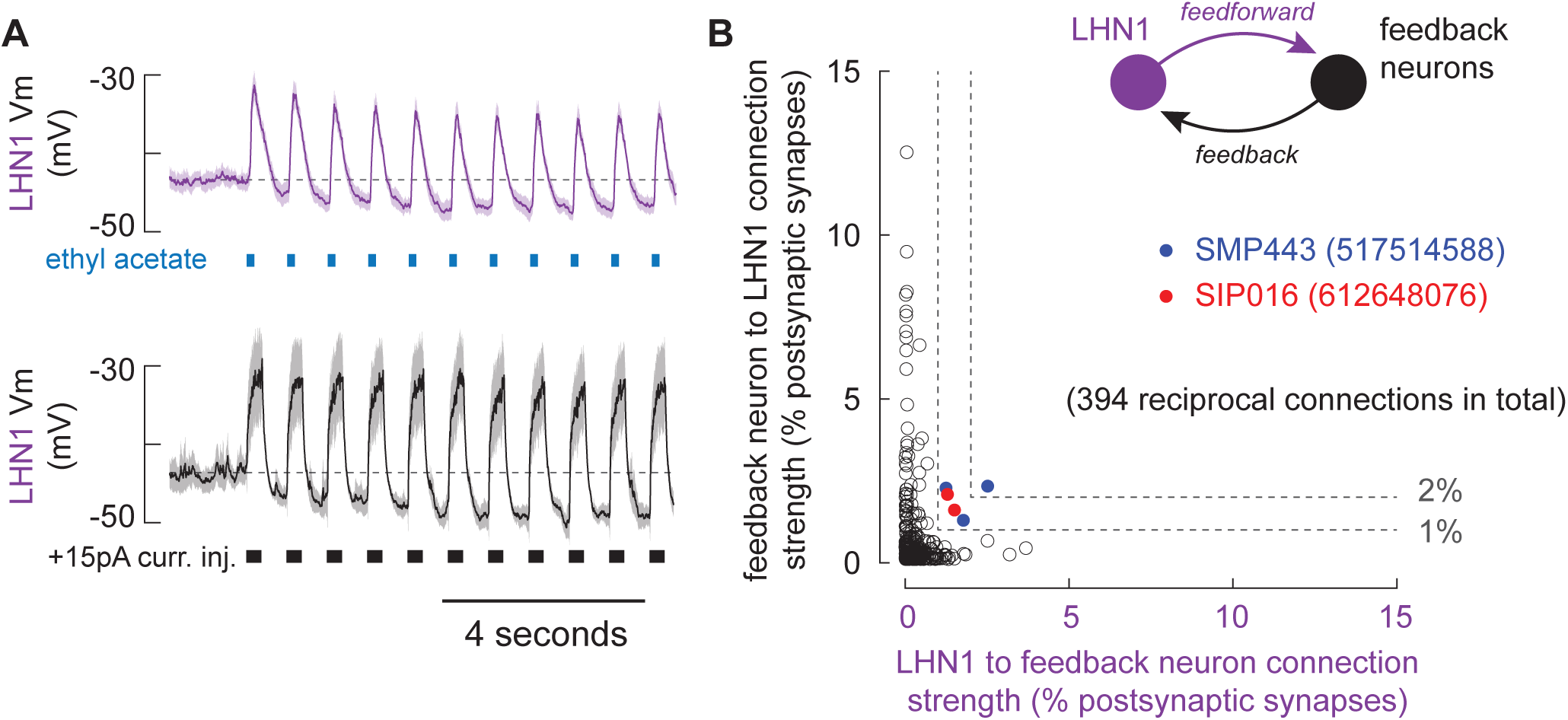
Odor-evoked afterhyperpolarizations LHN1 are likely of cellular origin. **(A)** Current injection is sufficient to evoke afterhyperpolarizations. Top: Mean (± s.e.m.) LHN1 membrane potential in response to 1.25 Hz ethyl acetate pulses (repeated from Figure 6A, top). Bottom: mean (± s.e.m.) response to +15pA current injection pulses for the same set of recordings. N = 5 recordings. **(B)** Comparison of connectivity for all 394 neurons in the hemibrain connectome that both receive input from LHN1 and send output to the same instance of LHN1. Each point denotes one reciprocal set of connections. Only two neurons get at least 1% of their input from an instance of LHN1 and also provide at least 1% of inputs to that same instance of LHN1, labeled in red and blue. There are multiple blue and red points in the scatter plot, since the same feedback neurons target multiple LHN1 instances. These two neurons have unknown neurotransmitters, but even if they were both GABAergic, they would constitute very minor reciprocal feedback pathways. Thus, the hyperpolarizations evident in panel (A) are likely of cellular origin.

